# Identification and classification of abundant RNA-binding proteins in the mouse lens and interactions of Carhsp1, Igf2bp1/ZBP1, and Ybx1 with crystallin and β-actin mRNAs

**DOI:** 10.1101/2025.01.10.632466

**Authors:** Danielle Rayêe, Dong-Woo Hwang, William K. Chang, Ilana N. Karp, Yilin Zhao, Teresa Bowman, Salil A. Lachke, Robert H. Singer, Carolina Eliscovich, Ales Cvekl

**Affiliations:** The Departments of Ophthalmology and Visual Sciences, Albert Einstein College of Medicine, Bronx, NY 10461, USA; Cell Biology, Albert Einstein College of Medicine, Bronx, NY 10461, USA; Developmental and Molecular Biology, Albert Einstein College of Medicine, Bronx, NY 10461, USA; Genetics, Albert Einstein College of Medicine, Bronx, NY 10461, USA; Department of Biological Sciences, University of Delaware, Newark, DE; Medicine (Hepatology), Albert Einstein College of Medicine, Bronx, NY 10461, USA

**Keywords:** Carhsp1, Ybx1, Igf2bp1/Zbp1, β-actin and crystallin mRNAs, lens, RNA-binding proteins

## Abstract

RNA-binding proteins (RBPs) are critical regulators of mRNAs controlling all processes such as RNA transcription, transport, localization, translation, mRNA:ncRNA interactions, and decay. Cellular differentiation is driven by tissue-specific and/or tissue-preferred expression of proteins needed for the optimal function of mature cells, tissues and organs. Lens fiber cell differentiation is marked by high levels of expression of crystallin genes encoding critical proteins for lens transparency and light refraction. Herein we performed proteomic and transcriptomic analyses of RBPs in differentiating mouse lenses to identify the most abundant RBPs and establish dynamic changes of their expression in differentiating lens. Expression analyses include highly abundant RBPs, including Carhsp1, Igf2bp1/ZBP1, Ybx1, Pabpc1, Ddx39, and Rbm38. Binding sites of Carhsp1, Ybx1, and Igf2bp1/ZBP1 were predicted in various crystallin and β-actin mRNAs. Immunoprecipitations using antibodies against Carhsp1, Igf2bp1/ZBP1, and Ybx1 confirmed their interactions with αA-, αB-, and γA-crystallin mRNAs. A combination of single molecule RNA FISH (smFISH) and immunofluorescence was used to probe *in vivo* interactions of these RBPs with αA-, αB-crystallin, and β-actin mRNAs in cytoplasm and nucleoplasm of cultured mouse lens epithelial cells. Together, these results open new avenues to perform comprehensive genetic, cell, and molecular biology studies of individual RBPs in the lens.

## Introduction

RNA-binding proteins (RBPs) represent a large and diverse category of proteins involved in RNA biology, such as the post-transcriptional and translational regulation of gene expression. The most common RNA-binding mechanisms employ one or more RNA-binding domains, such as the RNA recognition motif (RRM), a K homology (KH) domain, and cold-shock domain (CSD). Individual RBPs can interact with both single or double stranded RNA molecules (see Gerstberger et al., 2014; Darnell et al., 2018; Lindquist and Mertens, 2018). These RNA:RBP interactions are also influenced by RNA secondary structures as well as specific RNA and protein covalent modifications (see Barbiery and Kouzarides, 2020). Mutations in genes encoding RBPs disrupt cell functions and result in a range of developmental abnormalities in a variety of model organisms (see Musunuru and Darnell, 2001; Lukong et al., 2008; Gerstberger et al., 2014). Despite of major progress using high-throughput approaches to probe cellular and molecular mechanisms of RBPs (Licatalosi et al., 2008; Castello et al., 2012; Dominguez et al., 2018, Van Nostrand et al., 2020), the full range and spectrum of functions of the majority of RBPs remain unknown.

Studies of RNA biology and RBPs represent a highly dynamic and challenging field as the individual RNA levels change dramatically throughout the development and in response to external signals. The RNA molecules are localized both in the nucleus and cytoplasm; sometimes transported to specific subcellular regions such as synapses in neuronal cells (Wu et al., 2016). The RNAs are represented by both coding (mRNAs) and noncoding RNAs (i.e. rRNAs, tRNAs, snRNAs, lncRNAs, and miRNAs). The individual RNA molecules are always thought to be associated with a complement of RBPs (Gerstberger et al., 2014; Darnell et al., 2018). Regarding the mRNA biology, pre-mRNAs, and mRNAs interacts with a dynamic coat of RBPs that dictates their processing, localization, stability, translational activity, and ultimately their decay (Haimovich et al., 2013). The major fraction of RBP:mRNA interactions occur at both 5’- and 3’-UTR regions of individual mRNAs illuminating one of multiple dimensions of the transcriptome *cis*-regulatory code. Current studies of RBPs are accelerated by systematic analyses of their structure and function (Gerstberger et al., 2014), analyses of their RNA-binding mechanisms (Castello et al., 2012; Dominguez et al., 2018; Jolma et al., 2020) as well as studies of mRNA:miRNA interactions that also include specific RBPs (Treiber et al., 2017). Although multiple studies exist to identify RBPs that interact with ubiquitously expressed mRNA, such as β-actin, c-Fos, Cdkn1b/p27, and others, much less is known regarding mRNAs encoding tissue-specific genes expressed under the tight developmental control.

The ocular lens has served as an excellent model to study RBPs and their roles in tissue morphogenesis for over a decade (Lachke et al. 2011; Dash et al. 2015; Siddam et al. 2018; Dash et al. 2020; Shao et al., 2020; Lachke, 2022); however, a systematic analysis of the abundant RBPs expressed in the lens remains to be performed. Lens development originates from a cluster of lens progenitor cells that are generated in the anterior pre-placodal region of the vertebrate embryos to generate a symmetrical pair of the lens placodes (Gunhaga, 2011; Cvekl and Zhang, 2017). Through the reciprocal invagination between the lens placode and optic vesicle, a polarized lens vesicle is formed that gives rise on its anterior and posterior portions to the lens epithelium and highly elongated lens fiber cells, respectively. The mature lens fibers express very high quantities of crystallin proteins that fill the cytoplasmic space and are required for lens transparency and refraction (see Bassnett et al. 2011). While many insights into the transcriptional control of individual crystallin genes exist (see Cvekl et al., 2015), their post-transcriptional and translational regulations remain poorly understood (see Cvekl and Eliscovich, 2021).

The first evidence about the role of non-coding nucleotides in lens mRNAs is based on findings that *cis*-mutations in 5’-UTR of ferritin light chain (FTL, chromosome 19q13.3) mRNA known as iron response element (IRE) for iron regulatory protein (IRP), which inhibits ferritin translation, causes congenital nuclear cataract coupled with hyperferritinemia (Girelli et al., 1995; Beaumont et al., 1995). Likewise, a mutation in 5’-UTR of the SLC16A12 gene causes human age-related cataract (Zuercher et al., 2010). Importantly, mutations in a Tudor domain containing 7 (TDRD7), a component of cytoplasmic RNA granules, were linked to cataract and glaucoma (Lachke et al., 2011). Most recently, Celf1 (Siddam et al., 2018), Caprin2 (Dash et al., 2015), and Rmb24 (Grifone et al., 2018; Dash et al., 2020; Shao et al., 2020) were studied in mouse and/or zebrafish lens development and several novel molecular mechanisms of gene control within the lens were found.

The individual lens fibers can be viewed as large cellular factories to produce a small number of highly abundant crystallin proteins. Nevertheless, expression of other abundant lens proteins including intermediate beaded filament proteins Bfsp1 and Bfsp2 and membrane proteins Mip/aquaporin0, connexin 46/Gja3 and connexin 50/Gja8 does not reach crystallin levels. During lens embryogenesis, the length of individual fibers increases by a factor of 100-1,000 (Bassnett, 2005), requiring appropriate structural materials for the expanding cellular volumes. In mature lens fibers, crystallin protein concentration reaches ∼450 mg/ml levels (Bassnett et al. 2011) though transcription is limited by the highly controlled lens fiber cell denucleation process generating organelle-free zone in the central portion of the lens (“lens nucleus”) required for its transparency (Bassnett, 2009; Rowan et al. 2017; Brennan et al. 2021). Another important postnatal phenomenon is continuous lens growth and generation of new layers of outer lens fibers to expand the “lens cortex”. Lens transcriptional (Sun et al. 2015; Zhao et al., 2018), translational (see Berns and Bloemendal, 1974), and protein control (Liberek et al. 2008) systems are influenced by the denucleation process. To resolve these limitations, we propose that both transcriptional and translational factories evolved to both quantitatively and qualitatively “outperform” the average somatic cells that have a variety of options to adjust to new conditions. Identification of abundant RBPs in lens and those regulated during lens fiber cell differentiation is thus the first step towards a rational design of novel studies of lens post-transcriptional gene regulation.

In the present study, we aimed to analyze our recent lens proteomic (Zhao et al., 2019a) and transcriptomic (Zhao et al., 2018) data to identify and classify the most abundant lens RBPs throughout the lens fiber cells differentiation. We next selected a small group of representative RBPs for their expression analysis in the embryonic and newborn lenses. Finally, as a proof-of-principle we demonstrate *in vivo* association of Carhsp1 and Ybx1 with αA- and αB–crystallin, and Carhsp1, Igf2bp1/ZBP1 and Ybx1 with β-actin mRNAs in cultured mouse lens epithelial cell, respectively.

## Results

### Identification and classification of abundant mouse lens RBPs

The newborn (P0.5) mouse lens epithelium and fiber cell proteomes were initially analyzed for RNA-binding proteins present in top 525 proteins covering 90% of lens fiber cell proteins (Zhao et al. 2019a) yielding a total number of 62 RBPs. To extend this analysis, herein we analyzed top 2,000 proteins in both lens compartments using Celf1 (Siddam et al., 2018) ranked #1,746 in lens fiber cell proteome as a guidance for an extended cut within over 4,000 individual proteins found in each lens compartment (Zhao et al., 2019a). Each protein was examined using NCBI Gene database to support its classification as an established or a candidate RBP using two large-scale studies (Baltz et al., 2012; Castello et al., 2012). This analysis generated a total number of 283 and 133 proteins annotated as established and candidate RBPs, respectively (Fig. 1A). In addition, we found 73 individual ribosomal proteins and 49 translational initiation and elongation factors, with many of them directly interacting with various RNA molecules (Fig. 1A). These 283 proteins represent over 18% of 1,542 human RBPs identified earlier (Gerstberger et al., 2014). Within these proteins, a total number of 72 (28) were only found in the top 2,000 proteins in the lens epithelium (fibers), respectively (Fig. 1B). A similar distribution was found for candidate RBPs (Fig. 1B). Next, within the group of 283 RBPs, we also found 141 proteins classified as rRNA-, tRNA-, dsRNA-, lncRNA- and snoRNA-binding proteins as well as 49 proteins involved in miRNA biogenesis (Fig. 1C). Regarding candidate RBPs, we further classified them based on the subcellular localization between the nucleus and cytoplasm or both (Fig. 1D). The final step included detailed functional classification of 283 RBPs into eight categories as shown in Fig. 1E (see also supplemental data S1, Excel file). While the majority of proteins received only single classification, multiple roles are known for lens RBPs localized both in the nucleus and cytoplasm such as Celf1 (Siddam et al., 2018) and Tdrd7 (Lachke et al., 2011; Barnum et al. 2020) as well as for multifunctional Fxr1 (Darnell et al., 2009), Csde1 (Moore and von Lindern, 2018), and Msi2 RBPs (Sundar et al., 2021) studied in detail outside of the lens. We next focused on three functional groups, including mRNA stability, translation and degradation, as we predict that these RBPs are involved in post-transcriptional and translational control of the most abundant lens fiber cell specific proteins.

**Fig. 1.**
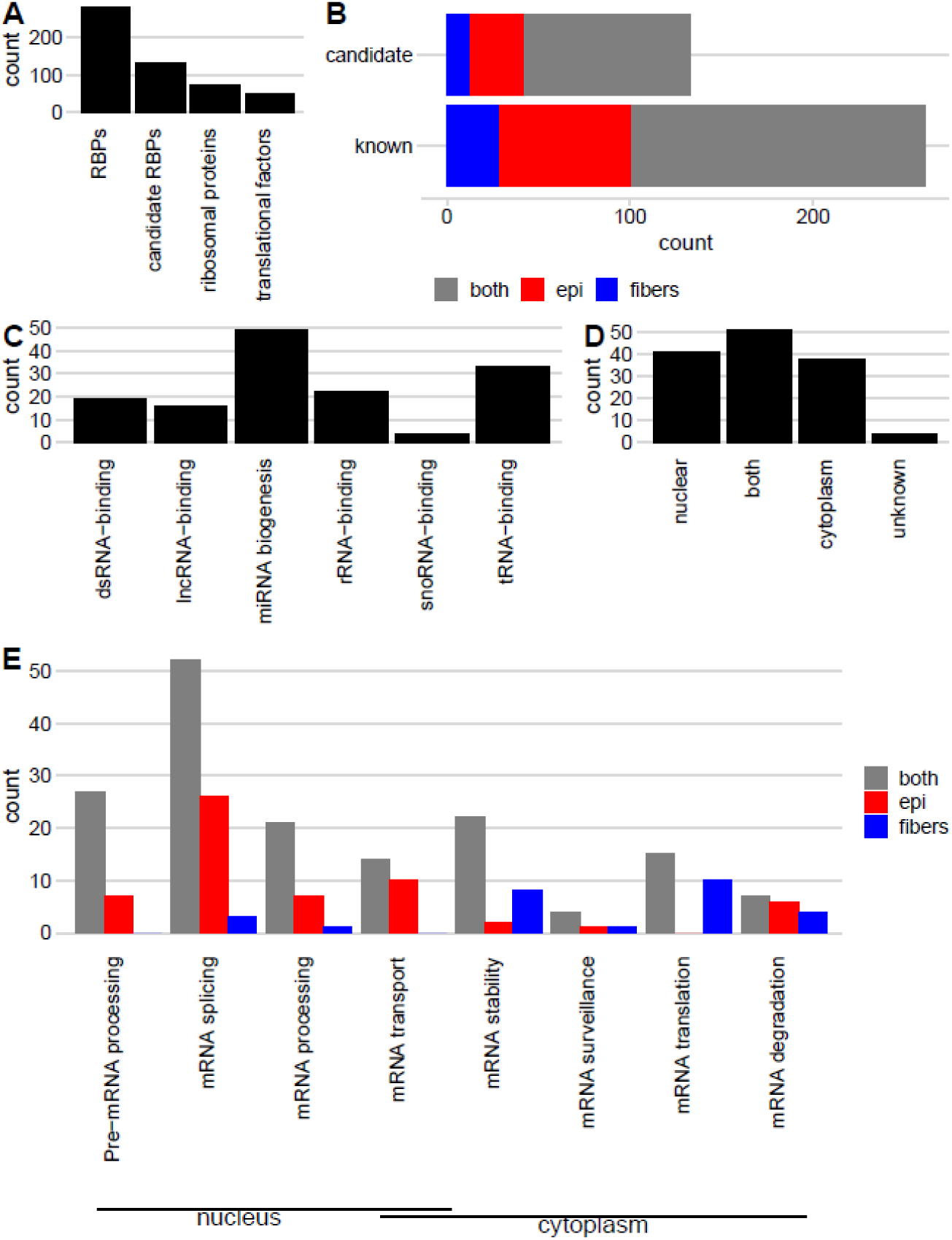
A summary of the most abundant mouse lens RBPs. **A.** Total number of RBPs, candidate RBPs, ribosomal proteins and translational factors in the analyzed set of proteins. **B.** Distribution of proteins found in lens epithelium, fiber and both compartments. **C.** Total number of RBPs based on their RNA-binding activity. **D.** Subcellular localization of candidate RBPs. **E.** General classification of 283 RBPs into eight categories, including their numbers in lens epithelium, fiber or and both compartments.

### Lens transcriptome: Differentially expressed genes encoding cytoplasmic RBPs

Our earlier studies using single cell analyses have shown that individual nascent crystallin mRNA transcription reaches the maximal outputs between E16.5 and P0.5 in lens fibers already undergoing preparation for the highly organized nuclear degradation (Limi et al., 2018; Limi et al., 2019). To identify differentially expressed RBPs during mouse lens differentiation, we thus analyzed bulk RNA-seq data of the mRNA stability, translation, and degradation groups using microdissected lens epithelium and lens fibers from embryonic stages E14.5, E16.5, E18.5, and newborn lens, P0.5 (Zhao et al. 2018). Lens differentiation cascade can be visualized through comparative data from E14.5 epithelium followed by E14.5 to P0.5 fiber cell expression data (Zhao et al 2019b). Within the group of 32 RBPs regulating cytoplasmic mRNA stability, heat maps revealed four main gene expression patterns, including moderate increase of transcripts in differentiating lens fibers, major up-regulation, moderate, and major down-regulation (Fig. 2A). To better visualize these data, the FPKM graphs using these five temporal/spatial stages are also shown in Fig. 2B. The most notable highly up-regulated mRNAs during lens fiber cell differentiation encode Carhsp1 > Park7 > Ybx1 > Tdrd7 > Pabpc1/Pabpc6 > Hspb1 > Rbm24/Rbm38 > Fxr1 > Ybx3 > Larp4 > Hsbp1 > Csde1; these genes are ordered based on their protein abundance in the lens fiber cells (Table 1). The most notable examples of down-regulated abundant mRNAs/proteins include Igf2bp3 > Igf2bp1/ZBP1 > Fmr1 (Fig. 2).

**Fig. 2.**
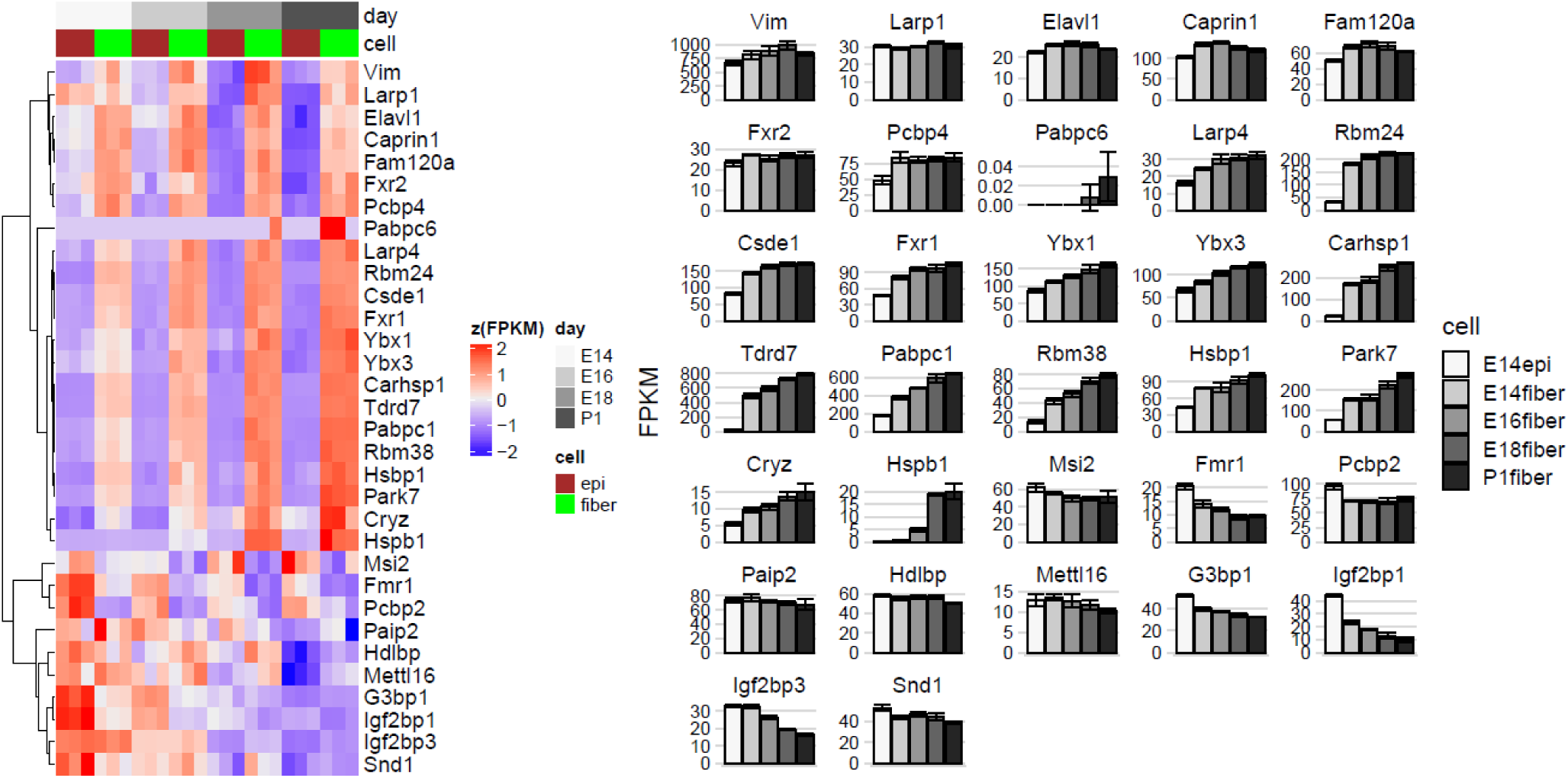
Expression profiles of 32 RBPs regulating mRNA cytoplasmic stability during lens differentiation. Barplots show FPKMs of the RBPs at E14.5 epithelium, E14.5 fibers, E16.5 fibers, E18.5 fibers, and P0.5 epithelium. Ybx1, Carhsp1, Rbm38, Pabpc1and Tdrd7 show increased expression whereas, Igf2bp1, Igf2bp3 and G3bp1 are downregulated throughout lens fiber cells development. Vimentin, Elavl1, Caprin1, Fxr2 and Pcbp2 display constant expression levels throughout fiber cells differentiation.

**Table 1.**
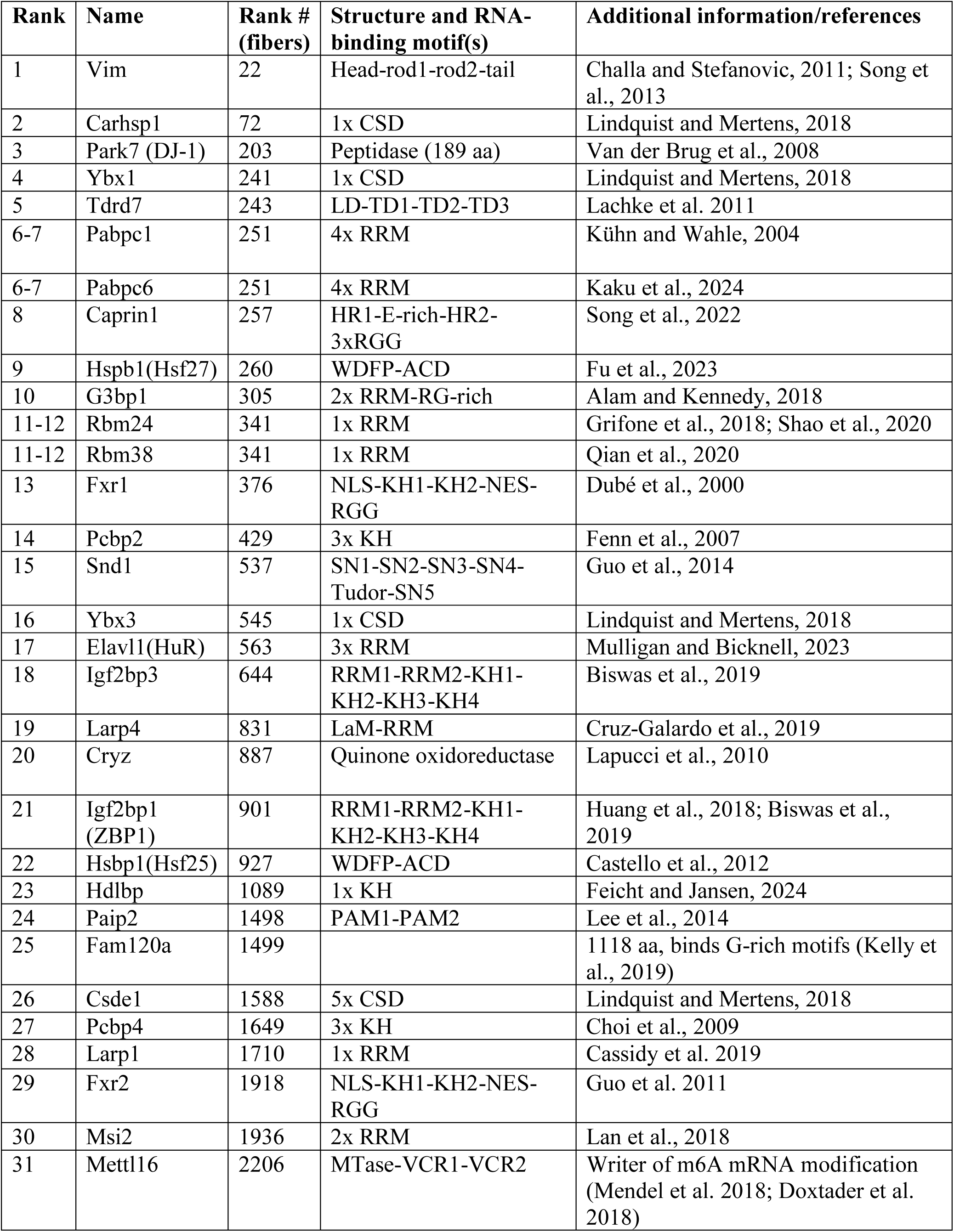
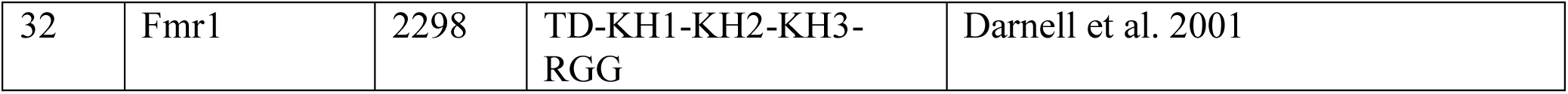
A summary of lens abundant RBP regulating mRNA stability.

Next, we analyzed 25 RBPs regulating translation (Fig. 3, Table 2). Their bulk RNA-seq data can be clustered into two main groups of transcripts, down- and up-regulated between epithelium and fibers (Fig. 3A). Eleven proteins with up-regulated mRNA include Caprin2 > Tdrd7 > Srp14 > Gspt1 > Larp4 > Etf1 > Csde1 > Paip1 > Celf1 > Atxn2 (Fig. 3B). Finally, within the group of 17 RBPs involved in mRNA degradation, bulk RNA-seq data identified moderate down- and up-regulation of 10 and 7 transcripts (Fig. 4). The up-regulated genes follow this protein ranking: Fxr1 > Gspt1 > Ddx6 > Dis3l2 > Lsm1 > Exosc4 > Trir (Fig. 4B, Table 3). Expression levels of five transcripts, encoding Rbm38, Rbm24, Carhsp1, Igf2bp1/Zbp1, Fxr1, Ybx1, Ybx3, G3bp1, and Igf2bp3, were independently validated by qRT-PCR (Supplemental Fig. S1).

**Fig. 3.**
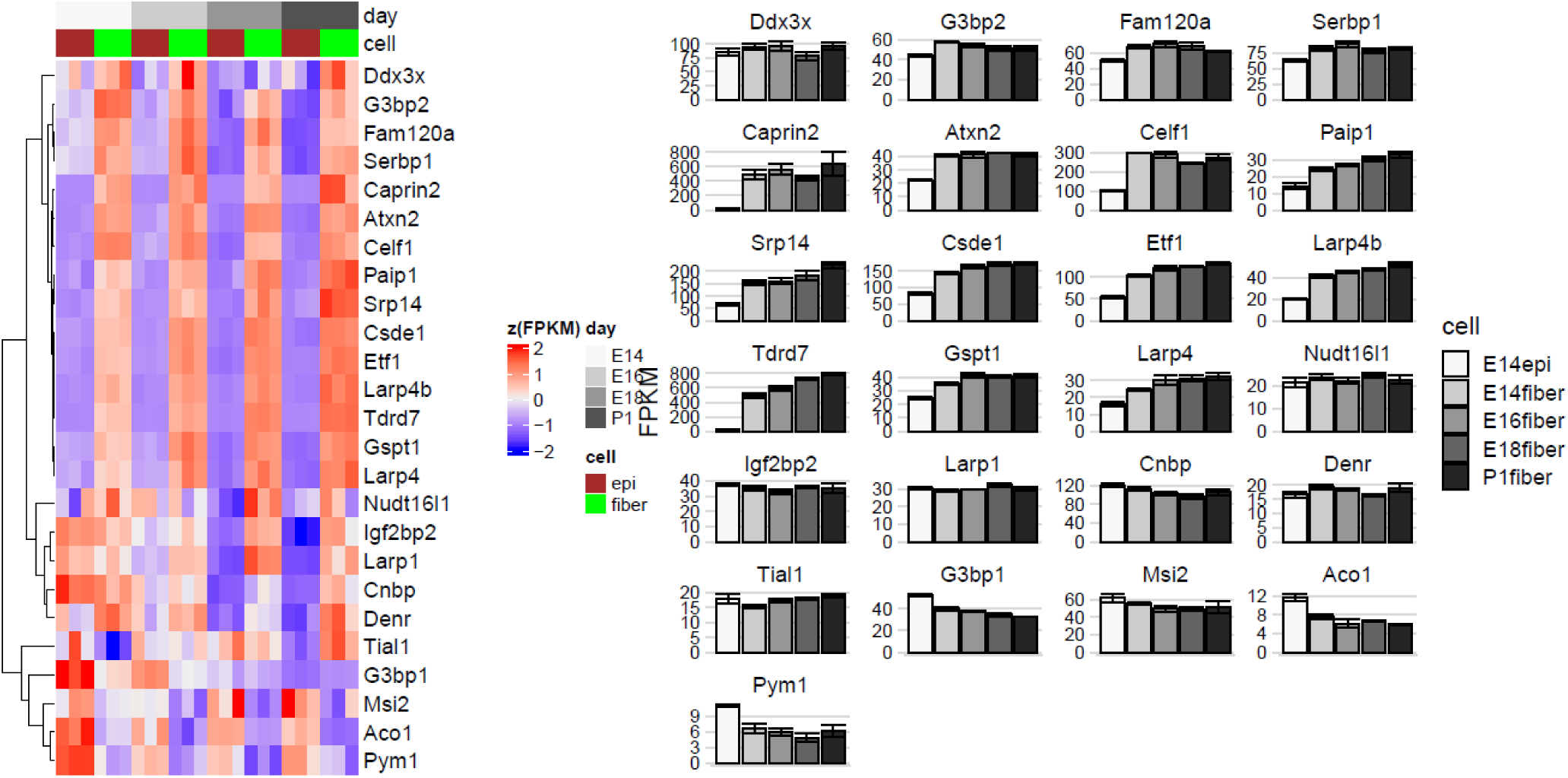
Expression profiles of 25 RBPs regulating translation during lens differentiation. Barplots show FPKMs of the RBPs at E14.5 epithelium, E14.5 fibers, E16.5 fibers, E18.5 fibers, and P0.5 epithelium. Tdrd7 is upregulated throughout fiber cells differentiation, whereas Celf1, Csde1, G3bp2 display constant expression levels.

**Fig. 4.**
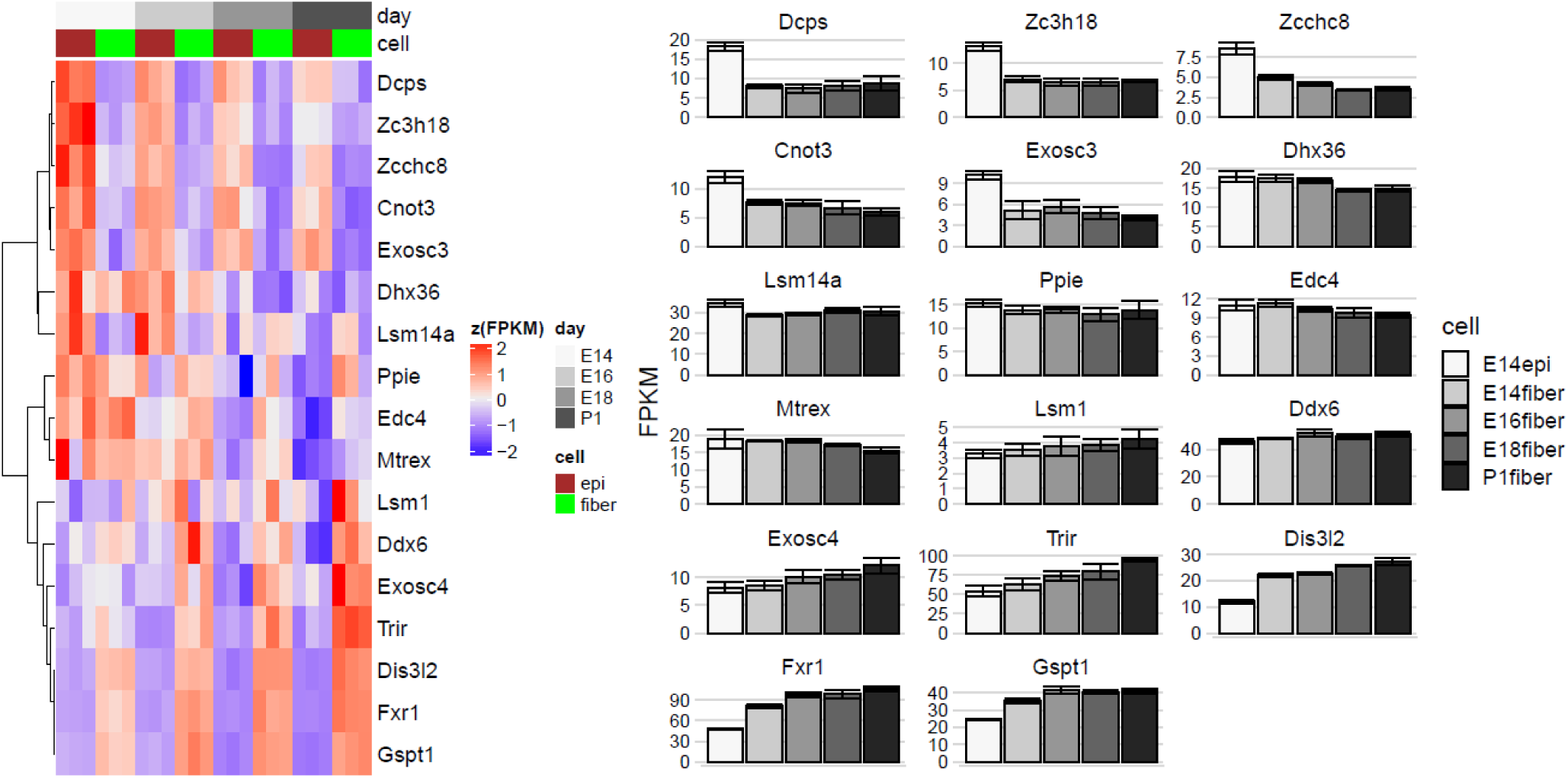
Expression profiles of 17 RBPs regulating mRNA degradation during lens differentiation. Barplots show FPKMs of the RBPs at E14.5 epithelium, E14.5 fibers, E16.5 fibers, E18.5 fibers, and P0.5 epithelium. Fxr1 is upregulated throughout fiber cell differentiation, whereas most of the other proteins involved in mRNA degradation are constantly expressed.

**Table 2.**
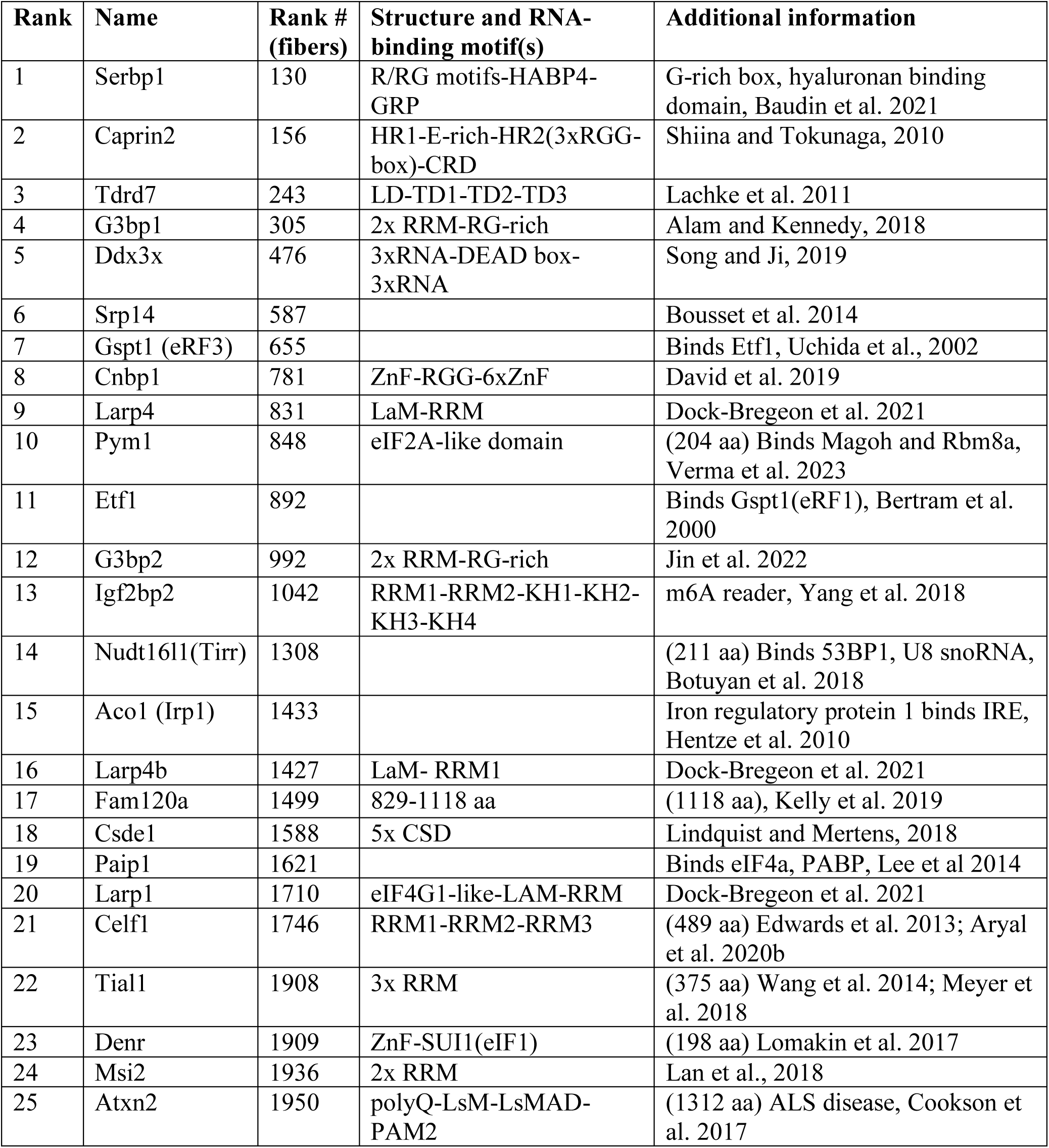
A summary of lens abundant RBP regulating mRNA translation.

| **Rank** | **Name** | **Rank # (fibers)** | **Structure and RNA-binding motif(s)** | **Additional information** |
| --- | --- | --- | --- | --- |
| 1 | Serbp1 | 130 | R/RG motifs-HABP4-GRP | G-rich box, hyaluronan binding domain, Baudin et al. 2021 |
| 2 | Caprin2 | 156 | HR1-E-rich-HR2(3xRGG-box)-CRD | Shiina and Tokunaga, 2010 |
| 3 | Tdrd7 | 243 | LD-TD1-TD2-TD3 | Lachke et al. 2011 |
| 4 | G3bp1 | 305 | 2x RRM-RG-rich | Alam and Kennedy, 2018 |
| 5 | Ddx3x | 476 | 3xRNA-DEAD box-3xRNA | Song and Ji, 2019 |
| 6 | Srp14 | 587 |  | Bousset et al. 2014 |
| 7 | Gspt1 (eRF3) | 655 |  | Binds Etf1, Uchida et al., 2002 |
| 8 | Cnbp1 | 781 | ZnF-RGG-6xZnF | David et al. 2019 |
| 9 | Larp4 | 831 | LaM-RRM | Dock-Bregeon et al. 2021 |
| 10 | Pym1 | 848 | eIF2A-like domain | (204 aa) Binds Magoh and Rbm8a, Verma et al. 2023 |
| 11 | Etf1 | 892 |  | Binds Gspt1(eRF1), Bertram et al. 2000 |
| 12 | G3bp2 | 992 | 2x RRM-RG-rich | Jin et al. 2022 |
| 13 | Igf2bp2 | 1042 | RRM1-RRM2-KH1-KH2-KH3-KH4 | m6A reader, Yang et al. 2018 |
| 14 | Nudt16l1(Tirr) | 1308 |  | (211 aa) Binds 53BP1, U8 snoRNA, Botuyan et al. 2018 |
| 15 | Aco1 (Irp1) | 1433 |  | Iron regulatory protein 1 binds IRE, Hentze et al. 2010 |
| 16 | Larp4b | 1427 | LaM- RRM1 | Dock-Bregeon et al. 2021 |
| 17 | Fam120a | 1499 | 829-1118 aa | (1118 aa), Kelly et al. 2019 |
| 18 | Csde1 | 1588 | 5x CSD | Lindquist and Mertens, 2018 |
| 19 | Paip1 | 1621 |  | Binds eIF4a, PABP, Lee et al 2014 |
| 20 | Larp1 | 1710 | eIF4G1-like-LAM-RRM | Dock-Bregeon et al. 2021 |
| 21 | Celf1 | 1746 | RRM1-RRM2-RRM3 | (489 aa) Edwards et al. 2013; Aryal et al. 2020b |
| 22 | Tial1 | 1908 | 3x RRM | (375 aa) Wang et al. 2014; Meyer et al. 2018 |
| 23 | Denr | 1909 | ZnF-SUI1(eIF1) | (198 aa) Lomakin et al. 2017 |
| 24 | Msi2 | 1936 | 2x RRM | Lan et al., 2018 |
| 25 | Atxn2 | 1950 | polyQ-LsM-LsMAD-PAM2 | (1312 aa) ALS disease, Cookson et al. 2017 |

**Table 3.**
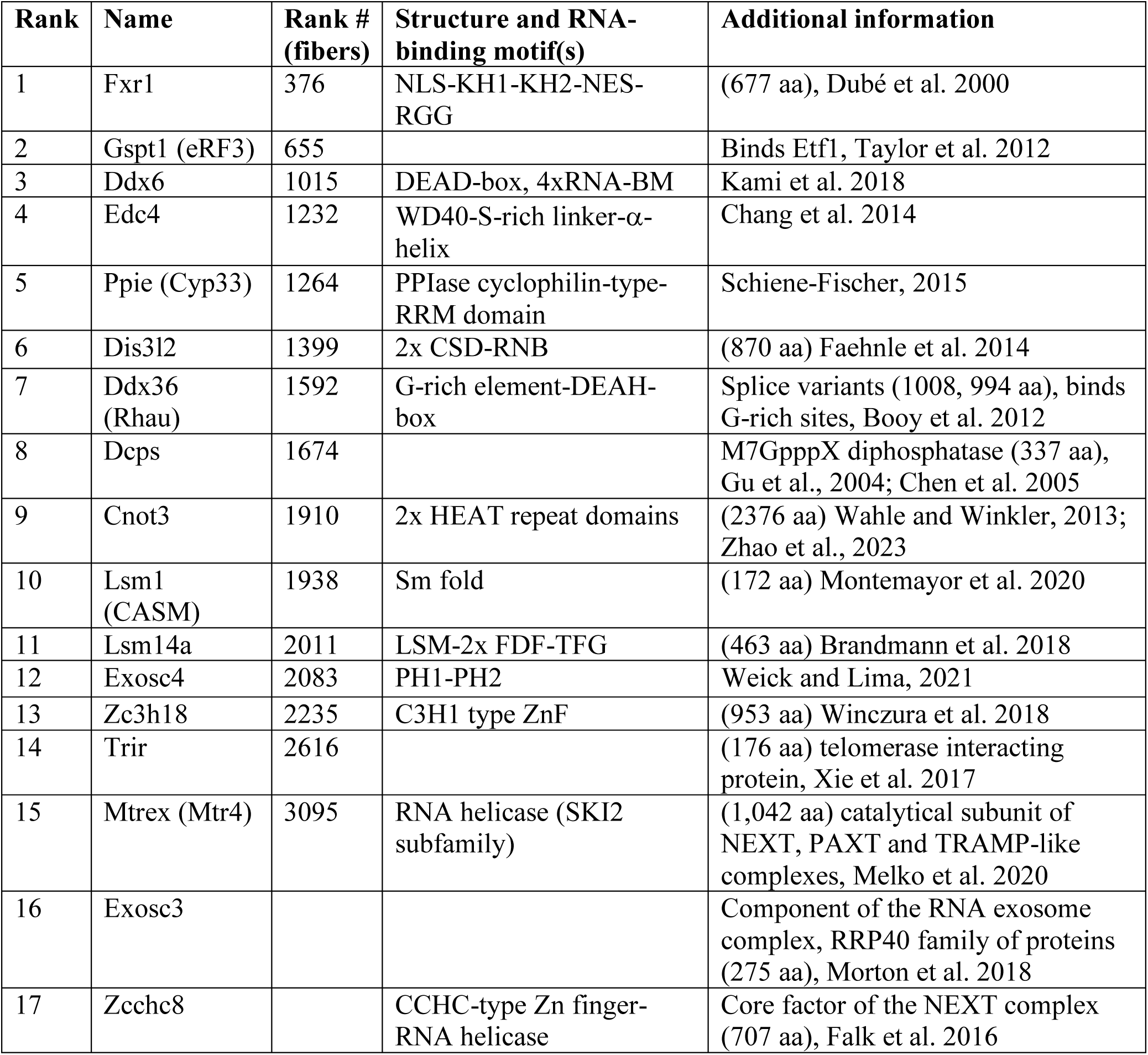
A summary of lens abundant RBP regulating mRNA degradation in the cytoplasm.

To gain additional information regarding expression of these RBPs at the RNA levels, we next analyzed expression of these RBPs both in early developmental (E10.5) and late adult mouse lens (2-months-old) using the iSyTE database (Kakrana et al., 2018). These data for “RNA stability” category show that Carhsp1 is highly expressed and enriched in lens from E12.5 to P56 (Fig. 5A,B) whereas Ybx1 is mostly expressed in the earliest primary lens fiber cells by E11-E12.5 (Fig. 5A). Tdrd7 is highly expressed in late fiber cells development, after E17.5 (Fig. 5A). These iSyTE data (Fig. 5B) also show that Carhsp1, Rbm24, Rbm38, and Tdrd7 are expressed much higher in the lens compared to other ocular tissues (Lachke et. 2012; Kakrana et al., 2018). Interestingly, Igf2bp1/ZBP1 is expressed throughout all stages of lens development with a marked peak of expression in late ocular lens at P56. Its expression in lens is also enriched when compared to other ocular tissues (Fig. 5B). Similar analyses were performed for “RNA translation” and “RNA degradation” (Supplementary Figures S4 and S5). Using data on postnatal lens epithelial and fiber cell gene expression (3, 6, and 24-months) highlights expression of these RBPs in cortical lens fiber cells containing nuclei. Taken together, this combination of proteomic and transcriptomic analyses demonstrates that lens fiber cell differentiation is marked by robust up-regulation of many cytoplasmic RBPs, and identifies individual proteins for follow up studies and design of RBP:mRNA interaction studies.

**Fig. 5.**
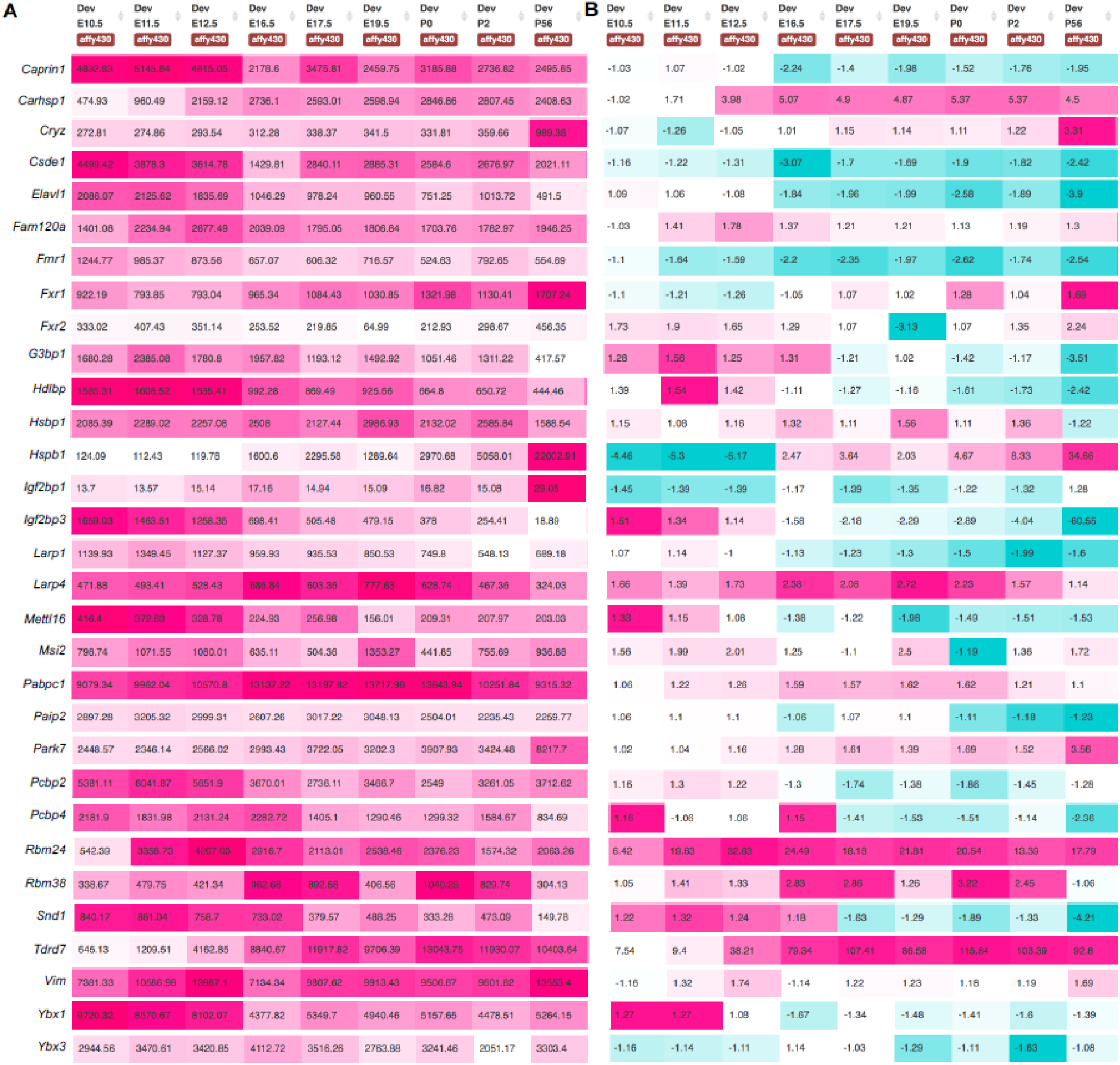
RBPs expression profile in early and late fiber cell differentiation. **A.** shows expression in fluorescence intensity units as reported by Affymetrix microarrays for different RBPs at various stages, embryonic day (E) 10.5 through 19.5 and at postnatal day (P) 0 (newborn) through P56. Row-specific heat-map reflects intensity of fluorescence signal (in turn reflecting expression) for individual RBPs at different stages. **B.** shows lens-enriched expression, given in fold-change, as compared to whole embryonic body (WB) reference data. Row-specific heat-map reflects extent of “lens-enriched” expression for individual RBPs at different stages.

### Localization of selected RBPs in the lens

To further extend our data, expressions of selected RBPs were analyzed in the developing mouse eyes with the focus on cytoplasmic, nuclear, or both localizations. Apart from vimentin, an intermediate filament protein regulating cytoskeleton, interacting with dsRNA from 5’-UTR (Zhang et al., 2022), the single Y-box (cold-shock domain, CSD) containing Carhsp1 (Pfeiffer et al. 2011; Zhao et al., 2019a) is the most abundant RBP in the lens fibers (Table 1). Carhsp1 is highly expressed in the cytoplasm of mouse lens from E12.5 to P0.5, however, some intra nuclear localizations were also found (Fig. 6A-D). Ybx1 (Kohno et al., 2003) is the fourth most expressed RBP in the lens fiber cells (Table 1) and is mostly found in the lens cytoplasm with some intranuclear signal detected after E14.5 (Figs. 6E-H). Note, that Ybx1 also functions as sequence-specific DNA-binding nuclear transcription factor recognizing sequence 5’-AACATGGT-3’ (Dolfini and Mantovani, 2013). We have also included Igf2b1/ZBP1 (Ross et al. 1997; Oleynikov and Singer, 2003), ranked #21 in the lens fiber cell proteome (Table 1), in the cellular localization analysis through immunofluorescence, as this protein has been shown to associate with the β-actin 3’UTR mRNAs within the neuronal cells (Eliscovich et al., 2017). Indeed, our earlier data have shown high levels of nascent expression of β-actin just prior the lens fiber cell denucleation (Limi et al., 2018). As expected, Igf2bp1/ZBP1 displays abundant cytoplasmic signals in the developing mouse lens (Fig. 6I-L).

**Fig. 6.**
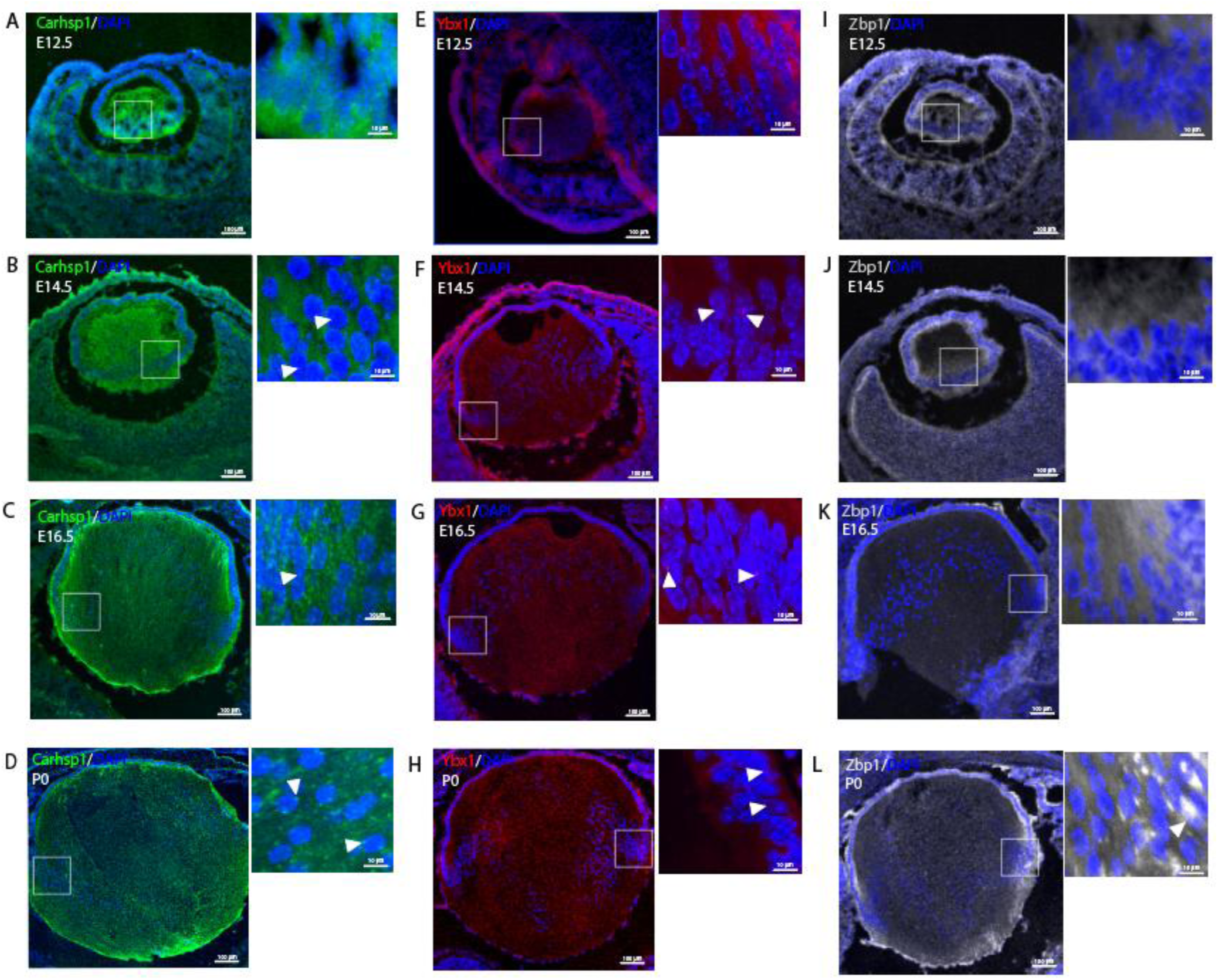
Most expressed RBPs localization in developing mouse lens tissue. **A-D** shows ocular lens from E12.5 to newborn (P0) immunolabeled with anti-Carhsp1 antibody (green). Carshsp1 stains predominantly fiber cell cytoplasm in all stages, with some intranuclear puncta seen after E14.5 (inserts, white arrowheads). **E-H.** shows ocular lens from E12.5 to P0 immunolabeled with antibody anti-Ybx1 (red). Ybx1 staining is predominantly cytoplasmic in all stages and intranuclear puncta staining seen from early to mature fiber cells in the transition zone area (inserts, white arrowheads). **I-L.** shows E12.5 to P0 lens immunolabeled with antibody anti-Zbp1 (grey). Zbp1 displays cytoplasmic signal only in all stages seen. Inserts location are indicated by white squares. Inserts represent 100X magnification.

### Identification of candidate RBP-binding sites within crystallin and β-actin mRNAs

To further investigate the interactions between crystallin mRNAs and candidate RBPs, individual crystallin mRNA sequences were analyzed for RBP-motifs using the Find Individual Motif Occurrences (FIMO) tool (Grant et al., 2011) (Fig. 7). Our analysis shows several binding sites for Carhsp1, Igf2bp1/ZBP1, and Ybx1 both in 3’UTR and 5’UTRs of crystallin mRNAs as well as within their coding sequences. We also used the bipartite RNA-binding motif 5’-CGGACN_10-25_(C/A)CA(C/U)-3’ found earlier in the previous studies of Igf2bp1/ZBP1 binding to the β-actin 3’-UTR and corresponding oligonucleotides (Chao et al., 2010; Patel et al. 2012; Eliscovich et al., 2017; Biswas et al. 2019). The αA-crystallin mRNA contains multiple Igf2bp1/ZBP1-binding sites as well as Carhsp1- and Ybx1-binding sites in its 3’UTR sequence. In contrast, the αB-crystallin mRNA displays multiple Carhsp1-binding sites, including a site located in the 5’UTR. We also found a binding site for Tardbp (fiber protein rank #317), encoding TDP-43 RNA-processing factor (Chen et al., 2019), overlapping Igf2bp1/ZBP1-binding site near the 5’UTR sequence of the αA-crystallin mRNA. In addition, we found an Elavl1 binding site near the Carhsp1-binding region in the αB-crystallin mRNA. It is possible that these additional RBPs have regulatory functions in association with our candidate RBPs. To further examine these mRNA:RBPs interactions, we used the γA-crystallin mRNA as an example of a very short crystallin sequence (about 700 nucleotides). The γA-crystallin mRNA displays binding sites for Carhsp1 located in the coding sequence near the 5’UTR region, as well as Ybx1 and Igf2bp1/ZBP1 binding sites. Importantly, the β-actin mRNA also contains multiple predicted binding sites for Carhsp1, Igf2bp1/ZBP1, and Ybx1. The bipartite Igf2bp1/ZBP1 sites were not found in any of 3’-UTRs of the αA- and αB-crystallins and two of these sites in the control β-actin mRNA are described in the legend to Fig. 7.

**Fig. 7.**
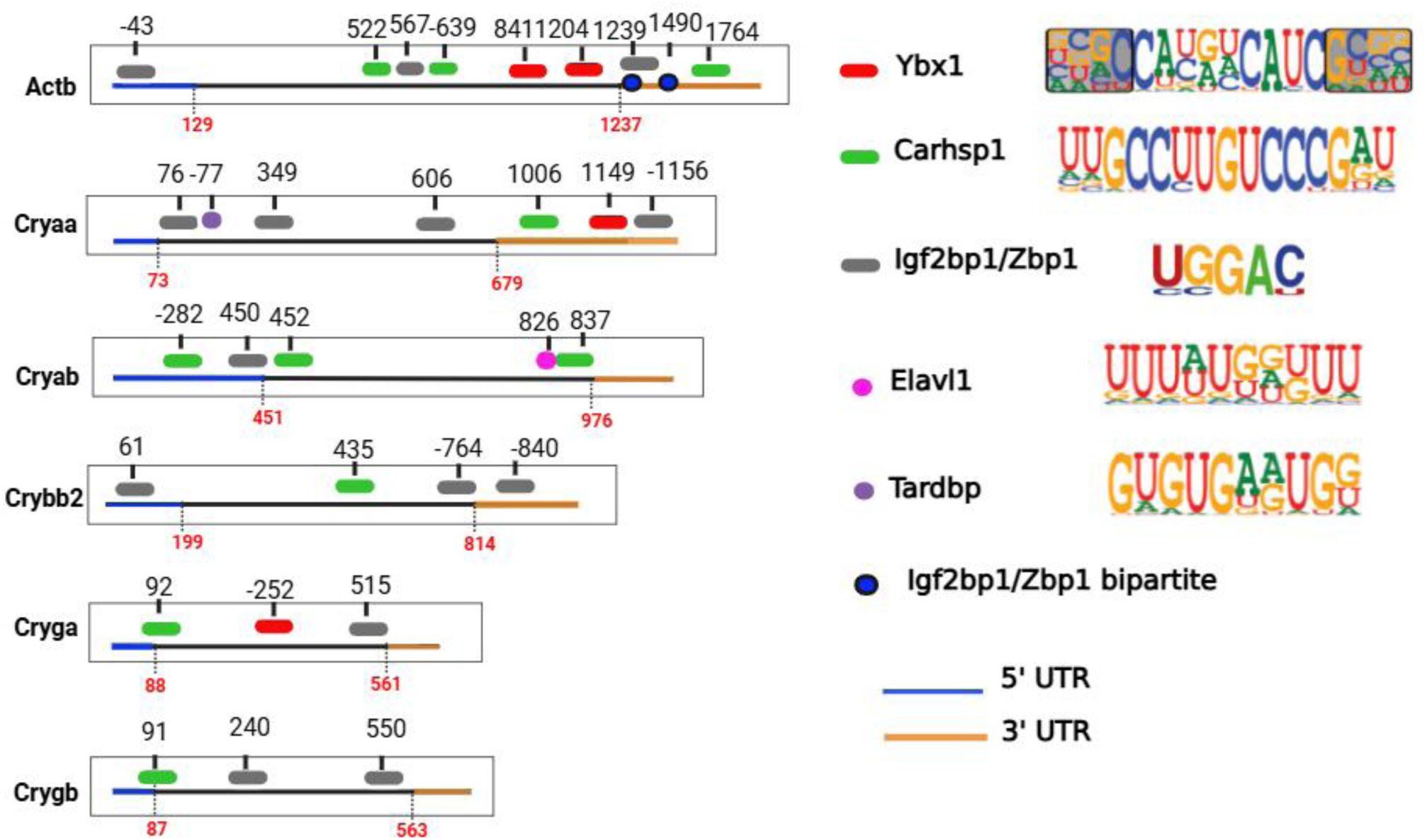
Metaplot of crystallin mRNAs and RBPs predicted interaction in mouse lens. Several crystallins display binding sites for RBPs represented in different colors. Ybx1 (red) is predicted to bind to the 3’UTR (shown in orange) sequence of αA-crystallin, 5’UTR (in blue) of αB-crystallin and the coding sequence (CDS) of γA-crystallin. β-actin displays two binding sites for Ybx1 in the CDS region. αA-crystallin displays one binding site for Carhsp1 (green) in the 3’UTR region, whereas αB-crystallin displays three binding sites: one in the 5’UTR region, one in the CDS and one flanking the 3’UTR sequence. Elavl1 (magenta) has a binding site overlapping Carhsp1 binding site near the 3’UTR sequence. Carhsp1 is predicted to bind γA-crystallin near the 5’UTR region, as well as β-actin in the CDS sequence (two binding sites) and in the 3’UTR. Several binding sites for Igf2bp1/Zbp1 (grey) were found in all sequences analyzed. Igf2bp1/Zbp1 is predicted to bind the 3’UTR region of αA-crystallin, the 5’UTR region of αB-crystallin and the CDS of γA-crystallin. αA-crystallin displays an overlapping binding site between Igf2bp1/Zbp1 and Tardbp (purple). β-actin mRNA displays three binding sites for Igf2bp1/Zbp1 being one in the 3’UTR region. The bipartite sequence (underlined) is located just one nucleotide downstream of the UGA stop codon: 5’-<u>CGGAC</u>TGTTACTGAGCTGCGTTTTAC<u>ACCC</u>-3’. The second bipartite site is located downstream: 5’-CUACAAAUGUGGCUGAGGAC-3’ (blue dots). Crystallin sequences are graphically represented by the black line and sequences size are shown by base pairs. End of 5’UTR and beginning of 3’UTR regions is shown by the base pair in red. Each RBP binding site is indicated by the base pair number equivalent to the beginning of the binding site sequence. Positive or negative numbers indicate either the positive or negative RNA strand where the binding site is located. RBP motifs are indicated on the bottom. 5’UTR is indicated in blue; 3’UTR in orange. αA-crystallin= Cryaa; αB-crystallin= Cryab; γ-crystallin= Cryga; β-actin= Actb.

### RNA-Immunoprecipitation (RIP) of crystallin mRNAs from the mouse lens extracts

To test our hypothesis that abundant lens RBPs interact with abundant crystallin mRNAs, we co-immunoprecipitated total RNAs extracted from newborn mouse lens whole cell extract using antibodies specific for Carhsp1, Ybx1 and Igf2bp1/ZBP1 proteins (Fig. 8A). We next generated cDNAs from the associated mRNAs and conducted a series of PCRs to detect specific enrichment of individual crystallin mRNAs (Fig. 8B). Both αA- and αB-crystallin mRNAs were enriched in Ybx1, Carhsp1, and Igf2bp1/Zbp1 immunocomplexes, whereas γA-crystallin mRNAs were enriched in Ybx1 and Igf2bp1/ZBP1 immunocomplexes only. Taken together, our findings show for the first time that Carhsp1, Ybx1, and Igf2bp1/ZBP1 may directly interact with highly abundant *Cryaa* and *Cryab* mRNAs.

**Fig 8.**
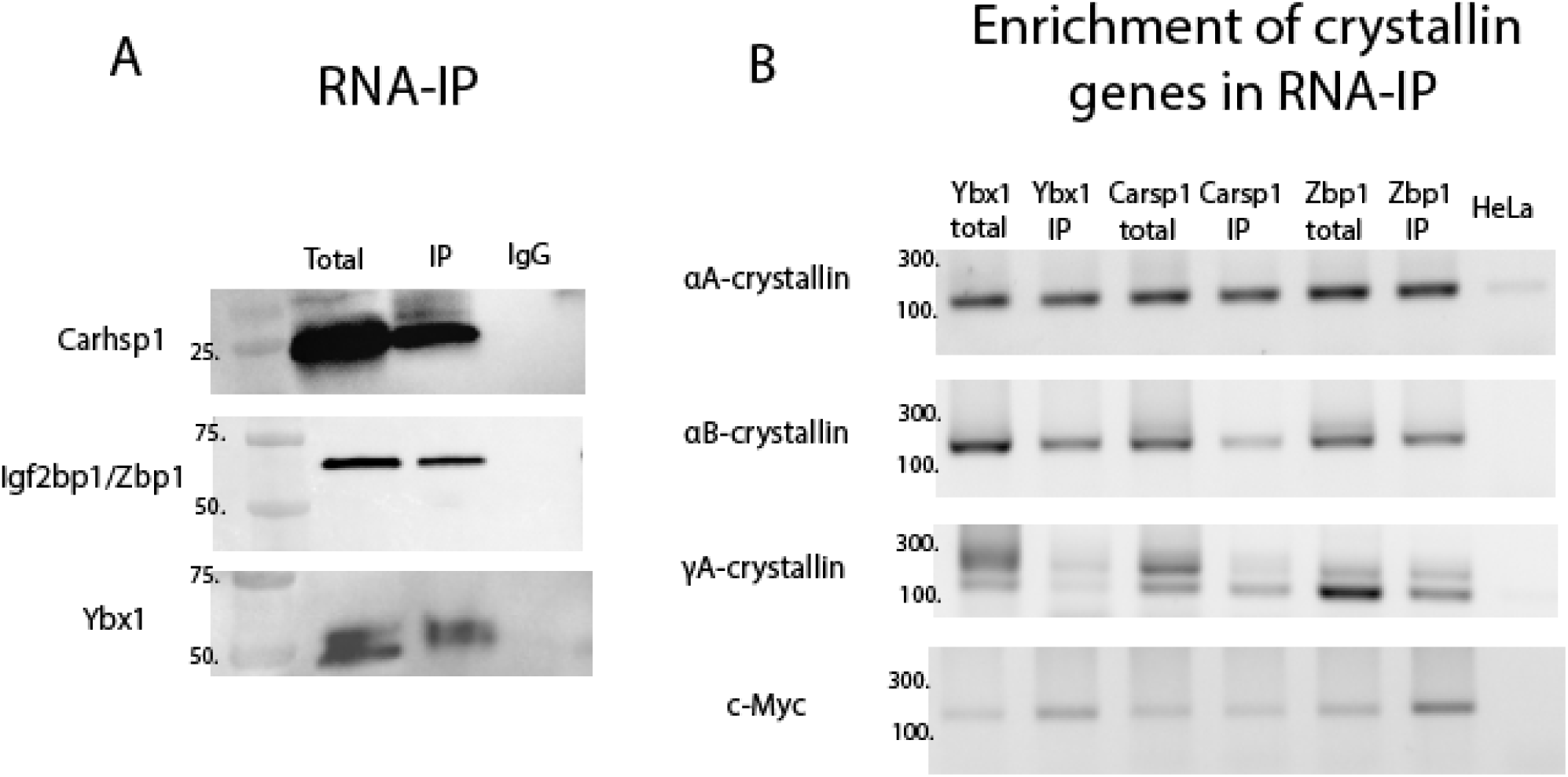
Enrichment of crystallin mRNAs in immunoprecipitated newborn lens. **A.** Newborn mouse lens extract was immunoprecipitated with antibody-coated beads for Carhsp1, Ybx1 and Zbp1. Western blot shows input and IP for each protein. **B.** RT-PCR of the immunoprecipitated RNAs shows enrichment of several crystallins. Ybx1-Immunoprecipitated RNAs show enrichment for αA and αB crystallins. Carhsp1-Imunnoprecipitated RNAs show enrichment for αA and αB crystallins. Zbp1-Immunoprecipitated RNAs show enrichment for αA, γA, and αB crystallins. Primers targeting c-Myc lens mRNA was included as a positive control to crystallin primers (bottom). Primers targeting HeLa mRNA were used as a negative control to crystallin primers (last lane).

### Single-molecule FISH (smFISH) of crystallins and β-actin mRNAs and their co-localization with RBPs

To further investigate the interactions of Carhsp1, Igf2bp1/ZBP1, and Ybx1, with lens crystallin mRNAs, we performed smFISH of αA and αB-crystallin mRNAs, in combination with immunofluorescence of Carhsp1, Ybx1, and Igf2bp1/ZBP1 proteins in cultured mouse lens-epithelial αTN4 cells (Figs. 9A, C, E). Individual mRNAs and RBPs were detected as discrete fluorescent spots at diffraction-level resolution in the cytoplasm and nucleus, enabling a detailed assessment of intermolecular spatial relations. Since subcellular localization of individual RBPs and mRNAs is subject to tight developmental regulation, we examined the extent of pairwise co-localization of mRNAs and RBPs in the nuclear and cytoplasmic compartments, by measuring the intermolecular distances between neighboring mRNAs and RBPs (Figs. 9B, D, F). Carhsp1 localized with αA- and αB-crystallin mRNAs mostly in the cytoplasm, considering total interactions found (Fig. 9B). Both Igf2bp1/ZBP1 and Ybx1 co-localized with αA- and αB-crystallins in both nuclear and cytoplasmic compartments (Fig. 9F). Interestingly, β-actin co-localized with all three RBPs in both cytoplasm and nucleus (Figs. 9B, D, F), as we have predicted above (Fig. 7). We have also found that β-actin mRNA is associated with Igf2bp1/Zbp1 in lens (Fig. 9D). Taken together, our results suggest that at least two crystallins mRNAs are associated with abundant lens RBPs such as Carhsp1, Ybx1 and Igf2bp1/Zbp1 within individual cells in mouse ocular lens (Fig. 9, Table 4).

**Fig. 9.**
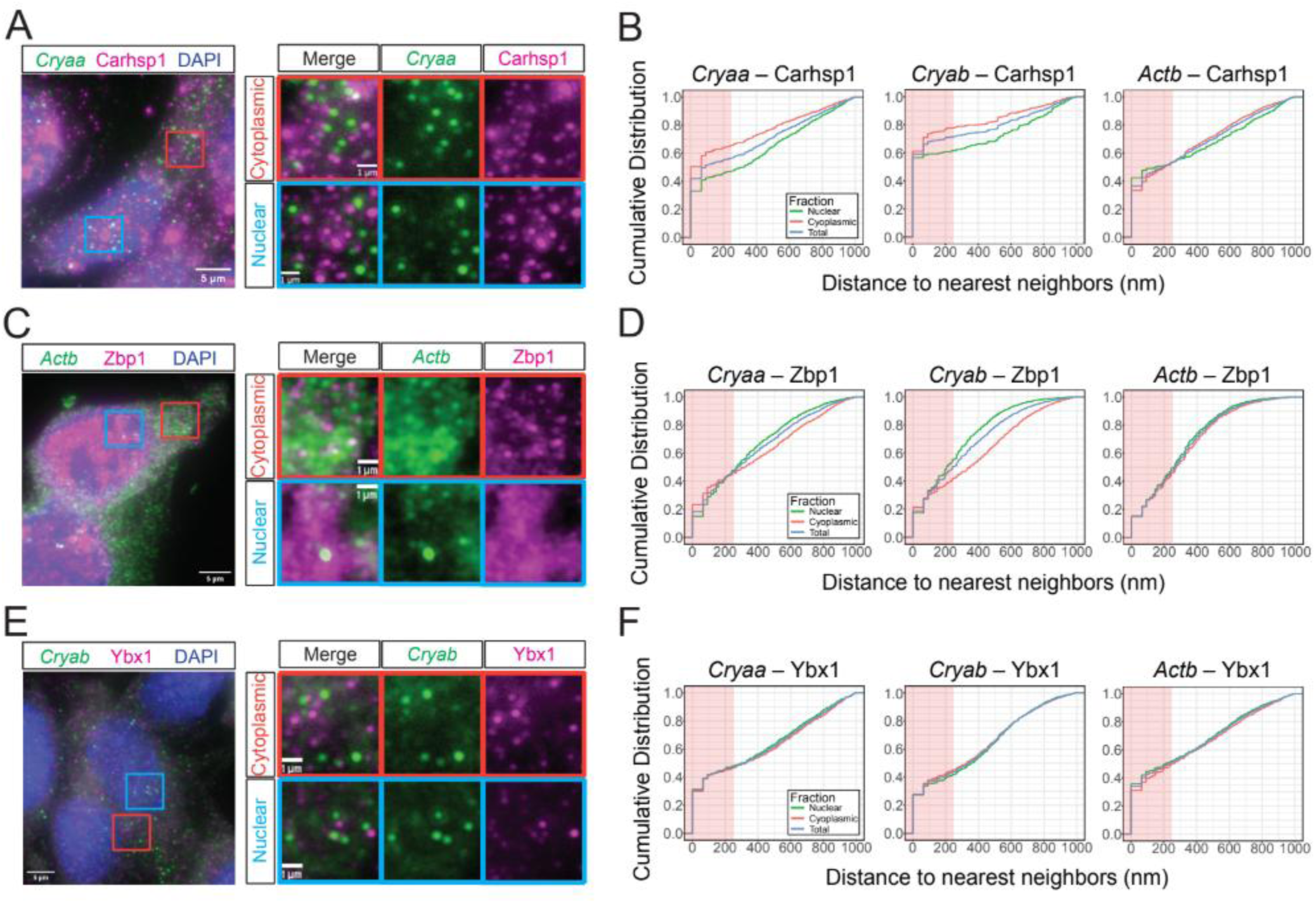
RBPs-mRNA interaction at single molecule level in lens-derived cell lines. Sub compartment-specific colocalization frequencies of mRNA-RBP pairs in lens-derived αTN4 cells. **A.** Representative results of single-molecule Fluorescent *in situ* Hybridization (smFISH), combined with immunofluorescence staining (IF), probing the αA-crystallin (Cryaa) mRNAs (green) and Carhsp1 (magenta) proteins in the nucleus (red inset) and cytoplasm (blue inset) in αTN4 cells cells. **B.** The fractions of colocalization between the Cryaa-Carhsp1, Cryab-Carhsp1 and Actb-Carhsp1 pairs in each subcompartment (i.e. nuclear and cytoplasmic) and cells (i.e. total) are shown in the cumulative distribution function plots as a function of distances to the nearest neighboring signals. The amber shade indicates the 250nm-colocalization distance threshold. **C.** smFISH representative results combined with IF, probing β-actin mRNAs (Actb, green) and Ig2bp1/Zbp1 in magenta. Proteins in the nucleus (red inset) and cytoplasm (blue inset) in αTN4 cells cells. **D.** The fractions of colocalization between the Cryaa-Zbp1, Cryab-Zbp1 and Actb-Zbp1 pairs in each subcompartment (i.e. nuclear and cytoplasmic) and cells (i.e. total) are shown in the cumulative distribution function plots as a function of distances to the nearest neighboring signals. The amber shade indicates the 250nm-colocalization distance threshold. **E.** smFISH representative results combined with IF, probing α-B crystallin (Cryab, green) and Ybx1 in magenta. Proteins in the nucleus (red inset) and cytoplasm (blue inset) in αTN4 cells cells. **F.** The fractions of colocalization between the Cryaa-Ybx1, Cryab-Ybx1 and Actb-Ybx1 pairs in each subcompartment (i.e. nuclear and cytoplasmic) and cells (i.e. total) are shown in the cumulative distribution function plots as a function of distances to the nearest neighboring signals. The amber shade indicates the 250nm-colocalization distance threshold. DAPI counterstaining in blue. αA-crystallin= Cryaa; αB-crystallin= Cryab; β-actin= Actb.

**Table 4.** Percentages of mRNAs:RBPs colocalized fractions in the nucleus, cytoplasm and total interactions found in lens-derived cell lines.

Subdepartment-specific colocalization frequencies of mRNA-RBP pairs
|  | Nuclear<br>colocalized fraction (%) | Cytoplasmic<br>colocalized fraction (%) | Total<br>colocalizaed fraction (%) |
| --- | --- | --- | --- |
| *Cryaa* – Carhsp1 | 39.78 | 53.24 | 45.95 |
| *Cryab* – Carhsp1 | 37.28 | 52.47 | 46.70 |
| *Actb* – Carhsp1 | 42.44 | 45.64 | 44.44 |
| *Cryaa* – Zbp1 | 44.70 | 38.84 | 42.21 |
| *Cryab* – Zbp1 | 54.82 | 39.32 | 48.87 |
| *Actb* – Zbp1 | 49.56 | 44.75 | 46.94 |
| *Cryaa* – Ybx1 | 38.84 | 35.26 | 36.25 |
| *Cryab* – Ybx1 | 38.71 | 41.27 | 40.20 |
| *Actb* – Ybx1 | 47.18 | 37.44 | 43.11 |

## Discussion

The mechanistic understanding of posttranscriptional and translational stages of individual gene expression in different cell types is driven by advanced studies of highly dynamic processes probing how individual RBPs interact with specific mRNA molecules, impact of these interactions on selective aspects of mRNA biology, and unique requirements of the individual cells depending on their cell-type identity and within the context of terminal differentiation. Multiple experimental challenges exist in this field as only some RBPs are thought to recognize specific RNA-binding motifs and RNA-molecules can fold to complex 3D-structures (Jolma et al., 2020). These proteins are mostly ubiquitously expressed, and their functional studies thus require gene loss-of-function studies in various cell types and different developmental stages (see Gertsberger et al., 2014). The present experimental model is mouse lens fiber cell terminal differentiation, and its unique feature is production of 16 individual crystallin proteins that reach levels that are only directly comparable with α- and β-globin gene expression in red blood cells (Sun et al., 2015). In both instances, degradation of nuclei is a major factor that further highlights operational demands for the entire cascade of gene expression (Rogerson et al., 2018).

Studies of gene expression at the level of transcription clearly show quantitative correlations between the repertoire of DNA-binding transcription factors and their subsequent roles in tissue-specific gene expression as shown in our earlier studies focused on the lens epithelium and lens fibers (Zhao et al., 2018; Zhao et al. 2019a). In the present study, we thus initially generated a census of the most abundant RBPs in mouse lens including their functional classifications followed by analyses of their expression dynamics at the bulk RNA levels during embryonic and early postnatal lens fiber cell differentiation (Figs. 1-4). Our initial summary demonstrates that most RBPs found here are present in both epithelial and fiber cells and are involved in mRNA splicing, processing and stability (Fig. 1E). In terms of their nuclear, cytoplasmic, and cytoplasmic/nuclear localization, these three categories include comparable numbers of RBPs (Fig. 1D). The earlier studied RBPs in the lens, Caprin2, Celf1, Rbm24, and Tdrd7, are ranked in lens epithelium proteome as #156, 1,746, 341, and 243, respectively. Earlier studies suggest interactions between Tdrd7 and βB3-crystallin mRNAs (Lachke et al. 2011, Barnum et al., 2020). Our protein census data using newborn lenses (mouse CD-1 strain) are supported also by recent proteomic studies of E14.5 whole lens (mouse C57Bl6 strain) proteome (Aryal et al., 2020a).

To identify candidate RBPs that bind crystallin mRNAs, we reasoned that the functional category “mRNA stabilization” is a logical starting point and the top five proteins are vimentin, Carhsp1, Park7, Ybx1, and Tdrd7 (see Table 1). Expression rank of Carhsp1 within all fiber cell proteins (proteomic rank #72) was followed by another CSD protein Ybx1 (#241) (see Lindquist and Mertens, 2018), and thus, both these proteins received experimental priority further enhanced by *in vivo* interactions between Ybx1 and β-actin mRNAs in neuronal cells (Eliscovich et al., 2017). This trio of proteins was extended by Pabpc1, DDX39, and Rbm38 analyzed in mouse eyes as shown in Figs. 6 and S2, respectively. The main conclusions from these studies are that both Carhsp1 and Ybx1 are highly up-regulated in differentiating lens fibers while reduction of expression of Igf2bp1/ZBP1 is notable at the bulk RNA-seq levels (Fig. 2). Analyses of iSyTE data further demonstrate that Carhsp1 is more expressed in lens compared to other ocular tissues while similar early score for Ybx1 is abolished after E12.5 (Fig. 5). Both Rbm24 and Rbm38 (Table 1) show upregulations of the encoding genes in lens fibers compared to the lens epithelium (Fig. 2). Our data agree with previous studies showing that Rbm24 is expressed in the lens epithelial and fiber cells, as well as the developing mouse inner ear (Grifone et al., 2018, Dash et al., 2020). Here we show that highly similar Rbm38 protein, linked to retinal differentiation in *Xenopus* (Dash et al., 2016) is lens-enriched, and upregulated by P1 (Figs. 2 and 5). Both Rbm38 and Rbm24 are upregulated in 3-month-old fiber cells (Fig. S3) (Dash et al., 2020). Rbm38 is found both in lens fiber cells cytoplasm and nucleoplasm (Fig. S2) suggesting its general role in stabilizing crystallin mRNAs. Importantly, studies of zebrafish Rbm24a identified its interactions with selected crystallin mRNA such as *cryaa* and its interactions with the cytoplasmic polyadenylation complex including Pabpc1a (Shao et al., 2020); however, the Rbm24a-binding sites remain to be identified.

Several genes encoding RBPs involved in RNA translation are up-regulated in newborn lens (Fig.3) as well as in aging lens such as Caprin2 and Celf1 (Fig. S3). Caprin2 is also significantly misexpressed following overexpression of FoxE3 in lens whereas expression of Celf1 is disrupted in the lens E2f1 knockout model (Table S4). The RBP-encoding genes engaged in RNA degradation show variable trends such as moderate down-regulation of Dcps, Cnot3 and Exosc3, and no significant changes (Lsp14a and Mtrex). In contrast, Fxr1 (fiber rank #376) that is up regulated in fiber cells from E14 to P1 and in 3-month-old mouse lens (Fig. 4,5). Fxr1 is known to regulate miRNAs involved in eye and cranial cartilage in *Xenopus* and its deletion leads to gross eye abnormalities (Gessert et al., 2010). Fxr1 is a highly conserved RBP, and our data shows for the first time that Fxr1 is continually expressed in mouse lens fibers. Taken together, these data suggest several novel RBPs in regulation of lens morphogenesis and cataract formation. Further studies are needed to better understand the mechanisms beyond their expression profiles.

To directly test our hypothesis that Carhsp1 and Ybx1 interact with crystallin mRNAs, our data using the FIMO tool (Grant et al., 2011) coupled with their recently established binding sites (Huang et al., 2018; Jolma et al., 2020) predict multiple Carhsp1- and Ybx1-sites in both αA- and αB-crystallin mRNAs (Fig. 7; Supplementary Table S3). We also predicted that αA-crystallin mRNAs displays several binding sites for Igf2bp1/ZBP1, one binding site for Carhsp1 and one binding site for Ybx1, both in the 3’UTR regulatory region. Subsequently, using RNA-IP (Fig. 8) we confirmed that Carhsp1 is associated with αA- and αB-crystallins mRNAs (Fig. 9D) to prompt the RNA hybridization-protein interaction studies by smFISH-IF. Earlier studies of Carhsp1 demonstrated its regulation of tumor necrosis factor alpha (TNF-α) mRNAs via its 3’UTR region to modulate the mRNA stability and decay by association with p-bodies; however, without any identification of the specific mRNA-binding sites (Pfeiffer et al., 2010).

The co-localization studies of mRNAs with individual RBPs serve as a powerful yet technically challenging tool to demonstrate *in vivo* this molecular mechanism (Eliscovich et al., 2017; Basyuk et al., 2021). Here we initially employed embryonic lens sections; however, given very strong signals of crystallin mRNAs, the subsequent analysis at the single-molecule resolution in combination with antibody staining was not technically possible. Nevertheless, using of cultured mouse lens epithelial cells used in previous studies of crystallin gene expression (Yang and Cvekl, 2005) made these experiments feasible. Here, we show for the first time Carhsp1 co-localizing with both αA- and αB-crystallin mRNAs within cultured lens epithelial cells suggesting an *in vivo* association of these RNA and protein molecules in the cytoplasm of lens cells. That might be crucial both to stabilize crystallin mRNAs to be used in multiple rounds of translation and to promote their accumulation in the expanding cytoplasmic volume of the elongating lens fiber cells. Ybx1 proteins also associate with αA- and αB-crystallin mRNAs in both the cytoplasm and nuclear compartments (Fig. 9, Table 4). Our FIMO motif analysis revealed a single Ybx1-binding site in the 3’UTR region of αA-crystallin mRNAs and another binding site within the protein coding sequence of the γA-crystallin mRNA. Importantly, we confirmed Ybx1 interaction with αA- and αB-crystallin mRNAs through RNA-IP and single-molecule fluorescence microscopy (Figs. 7 and 9). On the other hand, the motif analysis did not predict any Ybx1-binding site in the αB-crystallin mRNAs; however, both the immunoprecipitation and co-localization data support their mutual interactions. We propose that Ybx1 could be interacting with αB-crystallin mRNAs by yet unknown alternate binding motif or through association with another RBP, such as Elavl1 which displays an overlapping binding site near the Ybx1 binding site flanking the 3’UTR region (Fig. 7). It has already been shown that Ybx1 keeps mRNA stability by recruiting Elavl1, and both these RBPs form a complex (Chen et al., 2019). Here we show that Igf2bp1/ZBP1 co-localizes with αA- and αB-crystallins in both nuclear and cytoplasmic compartments (Figs. 7 and 9D). Finally, our data show the co-localization of Igf2bp1/Zbp1 and Ybx1 proteins to the β-actin mRNAs in cultured lens epithelial cells (Fig. 9, Table 4). Additional studies are underway to include non-RNA binding proteins as controls. Taken together, the present data open new research avenues for studies of selected RBPs and their roles in lens morphogenesis involving comprehensive genetic, cell, and molecular biology studies involving mapping of their interactions with mRNAs using CLIP-seq and HITS-CLIP to acquire panoramic views of the intricate complexity of mRNA:RBPs interactions (see Darnell, 2010).

## Materials and Methods

### Systemic analysis of RBPs expressed in the lens

Mouse proteomic data stored at the ProteomeXchangeConsortium via the PRIDE partner repository dataset identified PXD009639 were published earlier (Zhao et al. 2019a). To include all three previously studied lens RBPs, i.e. Caprin2, Celf1, and Tdrd7, the new cut off was set for the top 2,000 proteins in either lens epithelial or fiber cells. Three pairs of closely related proteins, including Ddx19a/Ddx19b, Pabpc1/Pabpc6, and Rbm24/Rbm38, list both proteins. rRNA-, tRNA-, dsRNA-, lncRNA- and snoRNA-binding and miRNA biogenesis functions were determined for each proteins using NCBI and published data (Shang et al., 2015; Treiber et al., 2017; Zhang et al., 2020; Jonas et al., 2020). Additional classification of RBPs included pre-mRNA processing, splicing, mRNA processing, mRNA transport, mRNA stability, mRNA surveillance, translational regulation, mRNA degradation, translational initiation, ribosome, and ncRNA binding (Gerstberger et al. 2014; Hentze et al. 2018; Dominguez et al. 2018; Van Nostrand et al. 2020). Nuclear and/or cytoplasmic or both localization of candidate RBPs were primarily determined using uniprot.org and proteinatlas.org web resources.

### Mice husbandry and tissue collection

Mice husbandry and tissue collection were conducted according to the approved protocol of the Albert Einstein College of Medicine Animal Institute Committee. All experiments were conducted following the ARVO statement for the use of animals in eye research. The day of confirmed vaginal plug was considered as the embryonic day E0.5. Eye tissues were harvested from embryonic mice at E12.5, E14.5, E16.5 and newborns (P0.5) CD1 mice. Embryonic eyes were collected after pregnant mice were euthanized by CO_2_ and mouse embryo heads were collected. Postnatal mice (P0.5) eyes were collected after euthanasia by CO_2_, followed by decapitation and eyeballs dissection. All tissues from embryonic and postnatal stages were fixed overnight in 4% paraformaldehyde at 4◦C followed by cryoprotection in 30% sucrose solution and embedded in optimal cutting temperature (OCT).

### Analysis of mouse lens RNA-seq and proteomic data and use of iSyTE

The RNA-seq datasets from lens epithelium and lens fibers contain four developmental stages, including E14.5, E16.5, E18.5, and P0.5 CD1 mice (Zhao et al. 2018). RBPs-encoding genes heatmaps and expression profile analysis were generated as we described earlier (Zhao et al., 2018). The iSYTE database (Lachke et al., 2012; Kakrana et al., 2018) was used to extend these analyses for both earlier E10.5-E11.5, and E12.5) and postnatal (P2, P56) stages on data obtained by expression microarrays. Further, “lens enriched expression” was obtained from lens vs. whole embryonic body (WB) reference dataset comparisons, as previously described (Lachke et al., 2012; Kakrana et al., 2018). iSyTE was also used to estimate differential expression of RBPs in different gene-specific perturbation mouse models as previously described (Kakrana et al., 2018, Choquet et al., 2021). In addition, publicly available RNA-seq data from isolated epithelium and fibers from 3, 6 and 24-months was used to estimate expression of RBPs in adult and aging adult stages (Whitson et al., 2017; Faranda et al., 2021).

### Analysis of crystallin mRNAs and microRNAs

The TargetScan database (v7.2) was used for crystallin mRNA seed-matching predictions (Lewis et al., 2005). The data on whole mouse lens ncRNAs, including E15 and P1 stages, were obtained from Khan et al., 2015. Additional information was gathered from miRNAs expressed in cultured rat lens cells (Wolf et al., 2013).

### FIMO motif analysis

Individual crystallin sequences were analyzed for RNA-binding protein (RBP) motifs using the Find Individual Motif Occurrences (FIMO) tool (Grant et al., 2011), with a significance threshold of p< 0.001. For all crystallins, full-length sequences (Supplementary Table S3), including the 5’ and 3’ untranslated regions (UTRs) of variant #1, were used. The presence of RBP motifs was confirmed by scanning the sequences for exact matches using a custom Python script. RBP motif sequences were derived from either an RBP pulldown assay targeting crystallin 3’ UTRs followed by mass spectrometry analysis, or from previously identified sequences from high-throughput studies (Jolma et al., 2020).

### Quantitative RT-PCR of RBPs in lens tissue

The RNAs were extracted from microdissected lens as we described earlier (Zhao et al., 2018). The RNAs were precipitated and their concentration measured by Nanodrop ND-2000 (Thermo Fisher, USA). Approximately 10 ng of cDNA was used for real-time PCR reaction mix following manufacturer’s protocol (SYBR Green PCR Master Mix, Sigma). mRNA fold change was calculated using Pcbp1 mRNA as housekeeping gene (O’Hagan et al., 2018).

### Antibodies and immunofluorescence

Lens sections were kept at room temperature for 1h followed by antigenic retrieval for 10 minutes at 90 ^0^C in Reveal Decloaker Solution (RV1000MMRTU, Biocare Medical). Sections were blocked in 10% NGS (50062Z, Thermo Fisher), 1% BSA (A4737, Sigma), 0.1% Triton (9036-19-5, Sigma) in PBS for 30 minutes at room temperature. Tissue sections were incubated at 4 ^0^C overnight with primary antibodies: anti-Carhsp1 (NBP1-31660, Novus Biologicals); anti-IMP1 (RN001M, MBL); anti-YBX1 (ab12148, Abcam). After overnight incubation, slides were washed in PBS and incubated for 2 hours at room temperature with secondary antibodies Goat anti-Rabbit IgG Alexa Fluor™ Plus 555 (A32732, Thermo) or Goat anti-Mouse IgG, Alexa Fluor™ 555 (A-21422, Thermo). Eye sections were washed in PBS, DAPI-stained for 5 minutes and mounted in ProLong™ Diamond Antifade Mountant (P36961, Thermo). Tissue images were acquired using the Leica SP8 Confocal at x10 or x63 oil-immersion objective.

### RIP using mouse lens

RNA immunoprecipitation from 200 mouse lenses was performed according to Jayaseelan et al. (2014). Briefly, lens tissue lysate was prepared using 1x Polysome Lysis Buffer [100mM KCl, 5 mM MgCl_2_ (M1028, Sigma), 10 mM HEPES, pH 7.0, 0.5 Nonidet P-40 (492016, Sigma Milipore)]. Dynabeads Protein A/G (100-02D; 100-04D, Invitrogen) were washed twice in NT-2 buffer [50 mM Tris pH 7.4, 150 mM NaCl, 1 mM MgCl_2,_ 0.05% Nonidet P-40] and incubated overnight at 4 degrees with 5µg of antibodies anti-Carhsp1 (A303-908A-T, Betyl Life Sciences), anti-YBX1 (ab12148, Abcam) and anti-IMP1 (RN001M, MBL). The magnetic beads were washed with NET-2 buffer [1x NT-2 buffer supplemented with 20 mM EDTA pH 8.0 (AM9260, Ambion), 1mM DTT and 200 units/ml RNAse OUT] and 100 µl of the cell lysate (20 mg/ml) was added. Before overnight incubation of the beads and lysate complex, 100 µl aliquot was taken to represent the ‘total’ RNA sample. The day after, the IPs were washed 6 times in NT-2 buffer, a 100 µl was taken and the standard western blot procedure was performed. IgG negative controls were added to each IP (3900S, Cell Signaling). The remaining IPs and total samples were digested in proteinase K buffer [1x NT-2 buffer, 1% SDS, 1.2 mg/ml Proteinase K (AM2546, Ambion)] and incubated at 55^0^ for 30 minutes. Phenol-chloroform RNA extraction was performed followed by cDNA synthesis following manufacturer’s instructions (18080051, SuperScript™ III First-Strand Synthesis System). RT-PCR using One-Taq Master Mix (M0482L, NEB) and primers for 3’UTR crystallin genes (Supplementary Table 2) was performed in total and IP RNAs.

### smFISH hybridization combined with immunofluorescence for RBPs

The murine lens epithelial cell line αTN4 (Yang and Cvekl, 2005) was cultured in Dulbecco’s Modified Eagle Medium containing 10% fetal bovine serum (Diecke and Beyer-Mears, 1997) and the smFISH protocol was performed according to our earlier studies with some modifications (Eliscovich et al., 2017). Briefly, cells washed three times in 1x PBSM (PBS supplemented with 5 mM MgCl_2_) and fixed in 4% paraformaldehyde in PBSM. After that, cells were washed in 1x PBSM followed by quenching in 0.1M Glycine/PBSM for 5 minutes at room temperature, followed by 1xPBSM wash. Coverslips were then incubated with permeabilization buffer (1x PBSM, 0.1% Triton) for 10 minutes at room temperature and washed in 1xPBSM two times for 10 minutes. Cells were then incubated with pre-hybridization buffer (30% deionized formamide in 2x SSC Buffer) for 30 minutes at 37◦C. αA and αB-crystallin probes mixture at 250 nM final concentration (IDT custom oligos, Oligostan, Supplementary Table 1) was added to the hybridization buffer [30% deionized formamide (AM 9342, Ambion), 1mg/ml E. coli tRNA (10109541001; Millipore Sigma), 10% dextran sulfate (D8906, SigmaAldrich), 0.2mg/ml Ultrapure BSA (10711454001, Roche), 2x Sodium Chloride-Sodium Citrate (SSC) buffer (AM9763, Invitrogen), 2 mM Ribonucleoside Vanadyl Complex (S1402S; New England Biolabs), UltraPure DEPC treated water (750023, Invitrogen)], and cells were incubated overnight at 37◦C. Cells were washed 2 times in 30% pre-hybridization buffer for 15 minutes, fixed in 4% paraformaldehyde in PBSM, washed two times and incubated in 10% pre-hybridization buffer for 30 minutes. The second set of readout probes (Supplementary Table 1) was diluted in 10% hybridization buffer, and cells were incubated for at least 3 hours at 37◦C. Coverslips were washed in 10% pre-hybridization buffer for 15 minutes, re-fixed in 4% paraformaldehyde, washed in PBS two times, and blocked in PBS 1% BSA for 30 minutes at room temperature. Cells were then incubated with primary antibodies anti-Carhsp1 (NBP1-31660, Novus Biologicals); anti-IMP1 (RN001M, MBL); anti-YBX1 (ab12148, Abcam) in blocking solution for 2 hours at room temperature. After that, coverslips were washed three times in PBS and incubated with secondary antibodies Goat anti-Rabbit IgG Alexa Fluor™ Plus 555 (A32732, Thermo) or Goat anti-Mouse IgG, Alexa Fluor™ 555 (A-21422, Thermo) for 30 minutes at room temperature. Coverslips were washed in PBS for 10 minutes, cell nuclei were counterstained with DAPI, and mounted in in ProLong™ Diamond Antifade Mountant (P36961, Thermo).

### smFISH imaging acquisition and analysis

Cells were imaged using a wide-field Olympus BX-63, supplied with a Super Apochromatic 100x1.35 NA objective, SOLA light engine (Lumencor), ORCA-R2 Digital Interline CCD Camera (C10600-10B; Hamamatsu) and filter sets DAPI-zero, Cy3-zero and Cy5-zero (Semrock). Total of 41 z-series per a field of view with the step size of 200 nm were acquired to cover 8 μm axial distance. A minimum of 50 cells per condition of randomly selected fields of views per experiment were acquired, using Metamorph software (Molecular Devices; RRID:SCR_002368) with the same acquisition settings and parameters. Initial inspection of the images was performed in Fiji (ImageJ). Selection of top 25 z-sections with the best sharp scores and subsequent segmentation of the nucleus and cell were performed, using the Big-FISH python package (Imbert et al., 2022). In Big-FISH, individual mRNA and RBP single molecule spots were identified as 3-d local maxima denoised and a Laplacian of Gaussian (LoG) filtered z-series images. Finally, the optimal intensity threshold was implemented based on the distribution of spot intensities, where a fast-decreasing population could be distinguished from a more stable population. Once the pixel coordinates of mRNA and RBP spots in the nucleus and cytoplasm were acquired, the distance between corresponding mRNA and RBP pairs were computed by the k-nearest neighbor algorithms (the k-value=5), using the SciKit-Learn python package (Pedregosa et al., 2011). Computed distances among the nearest neighbors in all fields of views across the mRNA-RBP pairs were then compiled, and the subcompartment-specific distributions across different pairs were visualized in the cumulative distribution function plots as a function of the distances to the nearest neighbors.

## Supporting information

Supplemental Files Rayee et al. 2025

## Acknowledgements

We thank Dr. Noura Ghazale for her helpful insights on the single-molecule imaging protocol.

## Abbreviations

CSD: cold-shock domain
FIMO: Find Individual Motif Occurrences
FISH: fluorescence in situ hybridization
KH: K homology
RBP: RNA-binding protein
RRM: RNA recognition motif
smFISH: single molecule RNA FISH
UTR: untranslated region

## Additional files

**Supplementary Fig S1. qRT-PCR of mRNA levels of most abundant RBPs in lens tissue throughout fiber cells differentiation.**

**Supplementary Fig. S2. Additional RBPs stainings in lens tissue.** A. Pabpc1 stainings at E14, E18 and P0. B. Ddx39 stainings at E14, E18 and P0. C. Rbm38 immunofluorescence staining in lens tissue at E12.5, E14.

**Supplementary Fig. S3. RNA-seq in adult and aged epithelial and fiber cells.**

**Supplementary Fig. S4. Microarray analysis of RBPs involved in degradation in early and late mouse lens.**

**Supplementary Fig. S5. Microarray analysis of RBPs involved in translation in early and late mouse lens.**

**Supplementary Table S1. Single-molecule RNA FISH hybridization probes**

**Supplementary Table S2. RT-PCR primers**

**Supplementary Table S3. RBP binding sites.**

**Supplementary Table S4. RBP encoding genes misexpressed in gene perturbation models.**

