## Supplemental Files Rayee et al. 2025 for "Identification and classification of abundant RNA-binding proteins in the mouse lens and interactions of Carhsp1, Igf2bp1/ZBP1, and Ybx1 with crystallin and β-actin mRNAs"

#### Supplementary Figure S1.

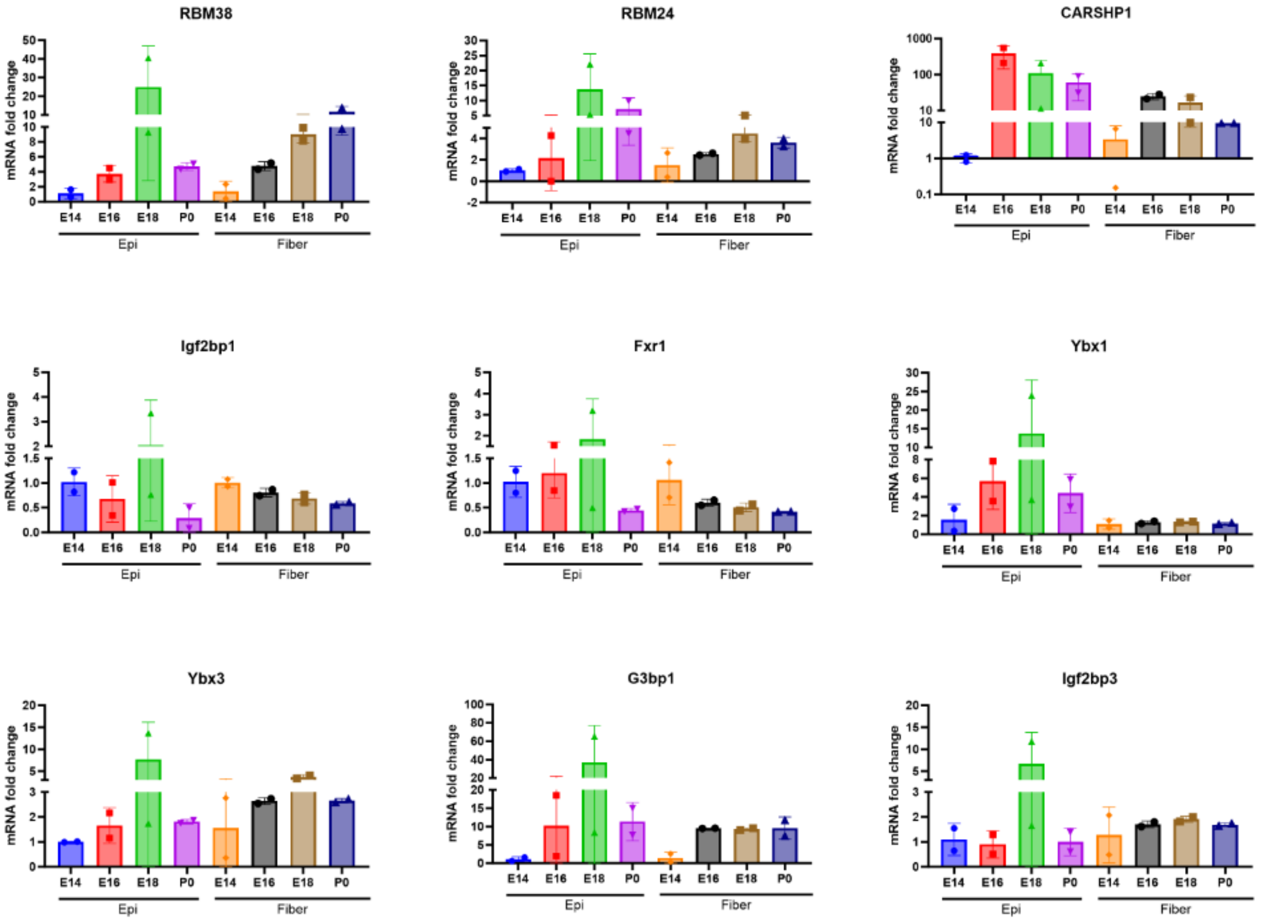

**Supplementary Figure S2.**

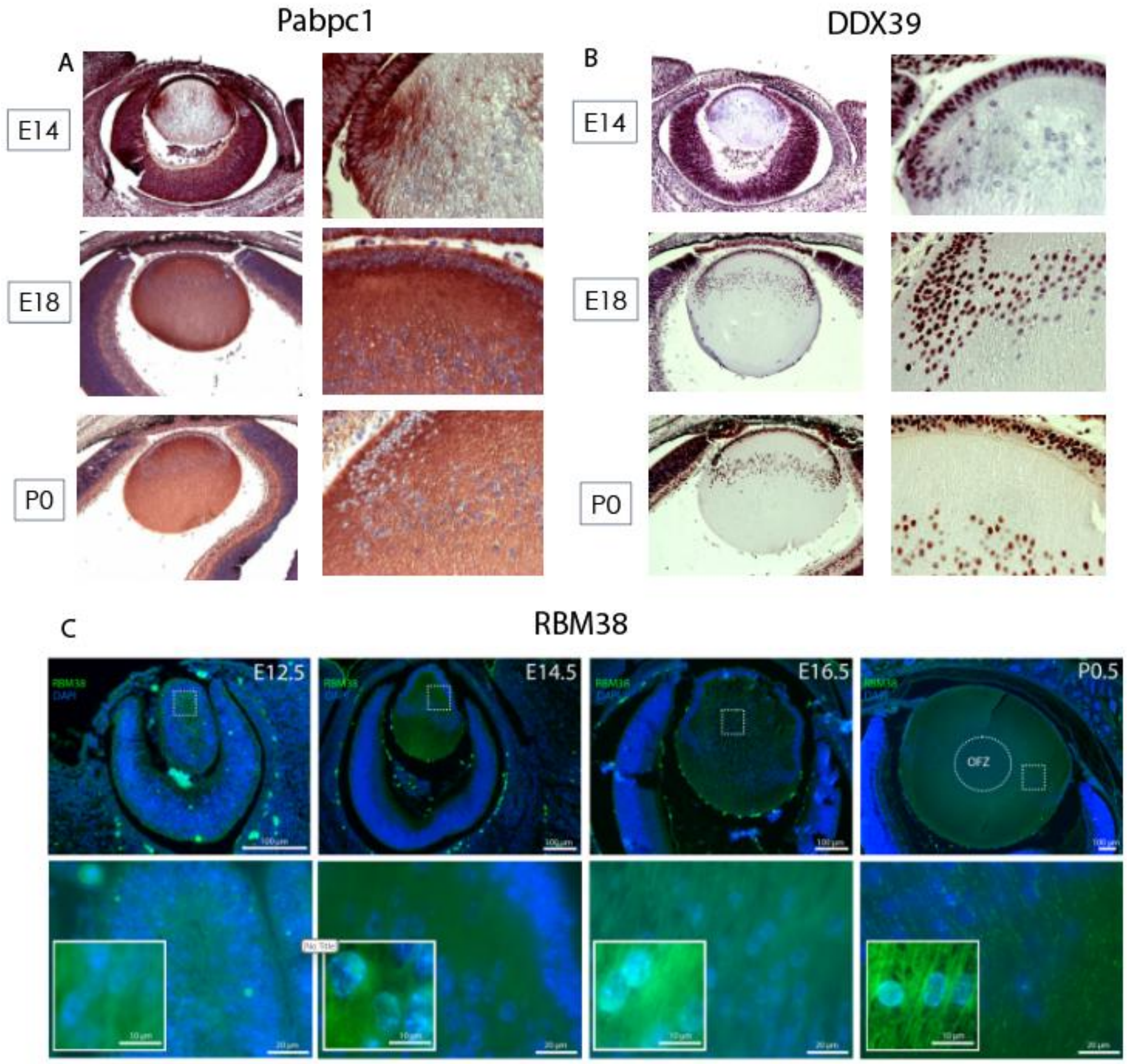

### Supplementary Figure S3.

A

| Gene | Epi 3 Mo | Fib 3 Mo | Epi 6 Mo | Fib 6 Mo | Epi 2 Yr | Fib 2 Yr |
|---|---|---|---|---|---|---|
| *Caprin1* | 21236.7 | 51740.0 | 19819.7 | 45856.9 | 18881.1 | 57449.9 |
| *Carhsp1* | 49042.4 | 172221.9 | 60158.4 | 270535.4 | 44710.4 | 169183.1 |
| *Cryzl1* | 5280.5 | 10744.3 | 11932.7 | 19564.6 | 4328.0 | 9711.1 |
| *Csde1* | 24825.5 | 43743.8 | 19186.2 | 39172.4 | 25941.6 | 53661.7 |
| *Elavl1* | 5975.3 | 5907.9 | 7537.0 | 8239.3 | 4771.7 | 6885.3 |
| *Fam120a* | 26223.4 | 32798.6 | 27086.5 | 31340.4 | 23181.5 | 31739.3 |
| *Fmr1* | 5664.7 | 6056.5 | 6585.1 | 6715.8 | 5874.4 | 8971.5 |
| *Fxr1* | 13509.4 | 44688.7 | 15082.9 | 43711.1 | 13137.0 | 36332.1 |
| *Fxr2* | 16486.4 | 61514.0 | 25351.1 | 98529.1 | 20498.1 | 75799.2 |
| *G3bp1* | 12320.8 | 9520.1 | 12911.8 | 9197.9 | 12204.3 | 11529.2 |
| *Hdlbp* | 26307.4 | 36508.2 | 23082.3 | 28692.3 | 30815.3 | 45212.3 |
| *Hsbp1* | 38415.9 | 102915.2 | 35828.3 | 72486.3 | 39210.6 | 73511.7 |
| *Hspb1* | 37680.3 | 4262380.8 | 8638.5 | 818238.5 | 20616.5 | 522656.8 |
| *Igf2bp1* | 1314.8 | 0.1 | 1471.7 | 27.1 | 1173.9 | 160.3 |
| *Larp1* | 25888.3 | 31011.5 | 19811.3 | 23296.8 | 27542.8 | 31510.4 |
| *Larp4b* | 11563.9 | 15719.8 | 13298.1 | 15928.1 | 10110.7 | 13602.0 |
| *Mettl16* | 2827.5 | 2511.4 | 2819.4 | 2585.8 | 2311.0 | 2584.1 |
| *Msi2* | 21438.9 | 29374.8 | 23173.2 | 31588.7 | 18532.0 | 24879.6 |
| *Pabpc1* | 55641.3 | 397453.4 | 40962.6 | 275743.7 | 57706.7 | 284515.0 |
| *Paip2* | 15226.4 | 28082.7 | 31316.3 | 44679.2 | 15396.9 | 24156.3 |
| *Park7* | 66320.7 | 422346.4 | 85567.2 | 504747.3 | 54801.9 | 234037.1 |
| *Pcbp2* | 27734.1 | 65718.2 | 36854.1 | 95081.3 | 30285.8 | 57081.4 |
| *Pcbp4* | 10524.2 | 13369.4 | 9796.7 | 22682.9 | 8185.9 | 14311.3 |
| *Rbm24* | 30518.6 | 112455.0 | 29921.5 | 118909.4 | 19620.2 | 68223.9 |
| *Rbm38* | 4326.0 | 28634.7 | 3433.3 | 19807.6 | 3388.9 | 16852.5 |
| *Snd1* | 6025.4 | 6104.6 | 7517.5 | 6467.9 | 6494.0 | 5959.5 |
| *Tdrd7* | 91918.9 | 918056.0 | 74264.0 | 830193.6 | 51817.0 | 432963.3 |
| *Vim* | 768710.6 | 1061098.8 | 674424.7 | 717474.6 | 771389.5 | 1059303.1 |
| *Ybx1* | 282709.7 | 835763.5 | 72367.0 | 225387.4 | 228043.5 | 652155.1 |
| *Ybx3* | 27930.8 | 91693.1 | 19011.8 | 72016.4 | 21543.1 | 59239.9 |

B

| Gene | Epi 3 Mo | Fib 3 Mo | Epi 6 Mo | Fib 6 Mo | Epi 2 Yr | Fib 2 Yr |
|---|---|---|---|---|---|---|
| *Aco1* | 5818.7 | 5162.4 | 8298.0 | 4300.3 | 6973.2 | 4794.4 |
| *Atxn2* | 9865.6562 | 10796.074 | 7209.298 | 7724.948 | 8694.9379 | 9347.5581 |
| *Caprin2* | 3966.365 | 21864.306 | 4464.7563 | 27131.429 | 2049.1387 | 9420.0871 |
| *Celf1* | 39463.2 | 82609.5 | 87475.8 | 182077.4 | 34914.5 | 80894.5 |
| *Cnbp* | 47334.0 | 76320.6 | 54150.5 | 88676.8 | 53229.8 | 81046.9 |
| *Csde1* | 24825.5 | 43743.8 | 19186.2 | 39172.4 | 25941.6 | 53661.7 |
| *Ddx3x* | 55252.01 | 51711.204 | 55193.426 | 58801.886 | 64530.92 | 71292.179 |
| *Denr* | 6515.9 | 9802.2 | 4903.8 | 9398.1 | 6758.6 | 11835.1 |
| *Etf1* | 52885.0 | 116831.4 | 62041.0 | 133021.5 | 43571.4 | 103733.2 |
| *Fam120a* | 26223.4 | 32798.6 | 27086.5 | 31340.4 | 23181.5 | 31739.3 |
| *G3bp1* | 12320.8 | 9520.1 | 12911.8 | 9197.9 | 12204.3 | 11529.2 |
| *G3bp2* | 16214.7 | 15756.0 | 21036.0 | 16335.9 | 17275.9 | 12139.7 |
| *Gspt1* | 17269.2 | 26201.6 | 14402.2 | 19928.1 | 14763.7 | 22262.8 |
| *Igf2bp2* | 2584.6 | 3605.4 | 987.0 | 672.1 | 145.2 | 0.1 |
| *Larp1* | 25888.3 | 31011.5 | 19811.3 | 23296.8 | 27542.8 | 31510.4 |
| *Larp4* | 5553.2255 | 18044.119 | 4443.4526 | 8170.5054 | 4991.1539 | 11612.925 |
| *Larp4b* | 11563.9 | 15719.8 | 13298.1 | 15928.1 | 10110.7 | 13602.0 |
| *Msi2* | 21438.9 | 29374.8 | 23173.2 | 31588.7 | 18532.0 | 24879.6 |
| *Nudt16l1* | 8567.1 | 4497.1 | 16181.3 | 13237.1 | 7865.8 | 7955.6 |
| *Paip1* | 14018.2 | 61419.8 | 19251.1 | 68874.5 | 13191.7 | 48672.0 |
| *Pym1* | 5677.1 | 4997.2 | 5071.8 | 3479.4 | 5645.1 | 2950.4 |
| *Serbp1* | 18062.7 | 53856.4 | 11873.7 | 36063.2 | 15220.1 | 39812.9 |
| *Srp14* | 68721.1 | 174764.3 | 47453.2 | 103112.1 | 59335.0 | 121725.8 |
| *Tdrd7* | 91918.9 | 918056.0 | 74264.0 | 830193.6 | 51817.0 | 432963.3 |
| *Tial1* | 7638.6 | 15645.6 | 19807.9 | 24995.9 | 6207.6 | 10110.1 |

C

| Gene | Epi 3 Mo | Fib 3 Mo | Epi 6 Mo | Fib 6 Mo | Epi 2 Yr | Fib 2 Yr |
|---|---|---|---|---|---|---|
| *Cnot3* | 7202.5 | 5957.8 | 5333.2 | 4260.3 | 7193.6 | 7219.7 |
| *Dcps* | 6137.4 | 6285.6 | 9318.1 | 9314.3 | 6347.2 | 5646.8 |
| *Ddx6* | 19456.6 | 19353.3 | 21537.1 | 12952.2 | 20666.9 | 25280.3 |
| *Dhx36* | 3898.5 | 3830.1 | 7860.5 | 5963.3 | 4089.3 | 4055.1 |
| *Dis3l2* | 2482.6 | 7202.3 | 1968.4 | 5536.0 | 2525.0 | 3312.2 |
| *Edc4* | 7367.6 | 6057.5 | 8956.9 | 7453.4 | 6948.6 | 6837.5 |
| *Exosc3* | 6417.3 | 2384.2 | 6451.3 | 4680.3 | 6059.5 | 5509.2 |
| *Exosc4* | 4360.0 | 6246.7 | 5826.2 | 11731.4 | 3101.1 | 8043.4 |
| *Fxr1* | 13509.4 | 44688.7 | 15082.9 | 43711.1 | 13137.0 | 36332.1 |
| *Gspt1* | 17269.2 | 26201.6 | 14402.2 | 19928.1 | 14763.7 | 22262.8 |
| *Lsm1* | 1801.7 | 1375.6 | 2593.5 | 2823.9 | 2055.3 | 1950.6 |
| *Lsm14a* | 11995.5 | 12810.2 | 14746.7 | 20622.4 | 11233.4 | 19869.6 |
| *Mtrex* | 6073.1 | 4028.9 | 7951.9 | 6512.0 | 5616.0 | 4994.9 |
| *Ppie* | 4599.1 | 4272.6 | 10110.2 | 9595.2 | 4728.0 | 4771.0 |
| *Trir* | 69874.7 | 435723.6 | 37398.3 | 239716.8 | 73070.5 | 213026.0 |
| *Zc3h18* | 5322.0 | 5085.6 | 3173.1 | 3803.7 | 5370.0 | 7099.6 |
| *Zcchc8* | 4497.1 | 2101.7 | 5756.7 | 3406.4 | 4005.9 | 4384.5 |

### Supplementary Figure S4.

A

| | Dev E10.5 affy430 | Dev E11.5 affy430 | Dev E12.5 affy430 | Dev E16.5 affy430 | Dev E17.5 affy430 | Dev E19.5 affy430 | Dev P0 affy430 | Dev P2 affy430 | Dev P56 affy430 |
|---|---|---|---|---|---|---|---|---|---|
| *Cnot3* | 972.79 | 876.56 | 843.55 | 571.08 | 414.61 | 350.76 | 374.41 | 435.7 | 328.39 |
| *Dcps* | 495.2 | 369.12 | 311.31 | 299.92 | 242.96 | 206.29 | 203.26 | 336.54 | 172.31 |
| *Dhx36* | 758.98 | 699.37 | 689.86 | 307.14 | 452.14 | 516.62 | 464.22 | 368.98 | 263.64 |
| *Dis3l2* | 113.98 | 108.68 | 135.78 | 195.26 | 179.55 | 152.1 | 184.01 | 177.54 | 79.06 |
| *Edc4* | 625.92 | 402.07 | 385.49 | 273.16 | 390.34 | 329.23 | 322.29 | 344.41 | 270.01 |
| *Exosc3* | 349.58 | 296.37 | 209.59 | 159.04 | 160.53 | 129.26 | 112.85 | 229.1 | 81.91 |
| *Exosc4* | 505.11 | 469.33 | 496.36 | 599.09 | 575.7 | 450.37 | 481.46 | 529.89 | 324.83 |
| *Fxr1* | 922.19 | 793.85 | 793.04 | 965.34 | 1084.43 | 1030.85 | 1321.98 | 1130.41 | 1707.24 |
| *Gspt1* | 1942.81 | 1909.39 | 2270.38 | 1691.74 | 1050.27 | 1249.22 | 879.54 | 1364.74 | 719.19 |
| *Lsm1* | 332.27 | 264.18 | 261.93 | 318.18 | 325.35 | 342.75 | 289.67 | 325.71 | 162.01 |
| *Lsm14a* | 1970.92 | 2138.43 | 1957.37 | 1538.6 | 1498.62 | 1527.52 | 1428.9 | 1247.08 | 1939.07 |
| *Mtrex* | 1219.54 | 835.65 | 666.8 | 809.34 | 663.46 | 657.83 | 589.17 | 532.79 | 467.56 |
| *Ppie* | 332.12 | 285.7 | 252.14 | 283.22 | 247.24 | 252.58 | 185.78 | 278.21 | 177.88 |
| *Trir* | 1599.19 | 1735.45 | 1508.53 | 1380.65 | 1545.87 | 1386.7 | 1654.21 | 1911.76 | 1894.53 |
| *Zc3h18* | 446.91 | 327.82 | 246.38 | 128.62 | 172.66 | 199.79 | 142.97 | 125.82 | 59.63 |
| *Zcchc8* | 577.06 | 533.33 | 381.23 | 284.29 | 213.61 | 358.71 | 168.99 | 229.75 | 140.91 |

B

| | Dev E10.5 affy430 | Dev E11.5 affy430 | Dev E12.5 affy430 | Dev E16.5 affy430 | Dev E17.5 affy430 | Dev E19.5 affy430 | Dev P0 affy430 | Dev P2 affy430 | Dev P56 affy430 |
|---|---|---|---|---|---|---|---|---|---|
| *Cnot3* | 1.58 | 1.92 | 1.61 | 1.09 | -1.31 | -1.54 | -1.45 | -1.16 | -1.65 |
| *Dcps* | 1.09 | -1.11 | -1.46 | -1.47 | -1.89 | -2.12 | -2.16 | -1.26 | -2.6 |
| *Dhx36* | 1.04 | -1.02 | -1.22 | -2.11 | -1.65 | -1.43 | -1.6 | -1.96 | -2.82 |
| *Dis3l2* | 1.24 | 1.32 | 1.55 | 2.23 | 1.94 | 1.69 | 2.06 | 2.01 | -1.14 |
| *Edc4* | 1.2 | -1.2 | -1.25 | -1.69 | -1.21 | -1.44 | -1.46 | -1.4 | -1.76 |
| *Exosc3* | -1.17 | -1.39 | -1.86 | -2.75 | -2.68 | -3.31 | -3.76 | -1.89 | -5.21 |
| *Exosc4* | 1.16 | 1.12 | 1.22 | 1.46 | 1.38 | 1.09 | 1.18 | 1.25 | -1.27 |
| *Fxr1* | -1.1 | -1.21 | -1.26 | -1.05 | 1.07 | 1.02 | 1.28 | 1.04 | 1.69 |
| *Gspt1* | 1.01 | 1.11 | 1.27 | -1.12 | -1.75 | -1.44 | -2.03 | -1.3 | -2.49 |
| *Lsm1* | -1.03 | -1.21 | -1.28 | -1.05 | -1.06 | -1 | -1.18 | -1.09 | -2.11 |
| *Lsm14a* | -1.05 | -1.06 | -1.07 | -1.23 | -1.31 | -1.28 | -1.37 | -1.55 | -1.01 |
| *Mtrex* | 1.16 | -1.18 | -1.29 | -1.19 | -1.46 | -1.48 | -1.63 | -1.97 | -2.11 |
| *Ppie* | 1.35 | 1.22 | 1.03 | 1.18 | 1.01 | 1.06 | -1.27 | 1.14 | -1.34 |
| *Trir* | 1.21 | 1.28 | 1.14 | 1.09 | 1.22 | 1.1 | 1.31 | 1.52 | 1.5 |
| *Zc3h18* | 1.57 | 1.36 | -1.14 | -2.09 | -1.71 | -1.44 | -2.01 | -2.25 | -4.88 |
| *Zcchc8* | 1.26 | 1.23 | -1.23 | -1.52 | -2.09 | -1.26 | -2.65 | -1.82 | -3.2 |

#### Supplementary Figure S5.

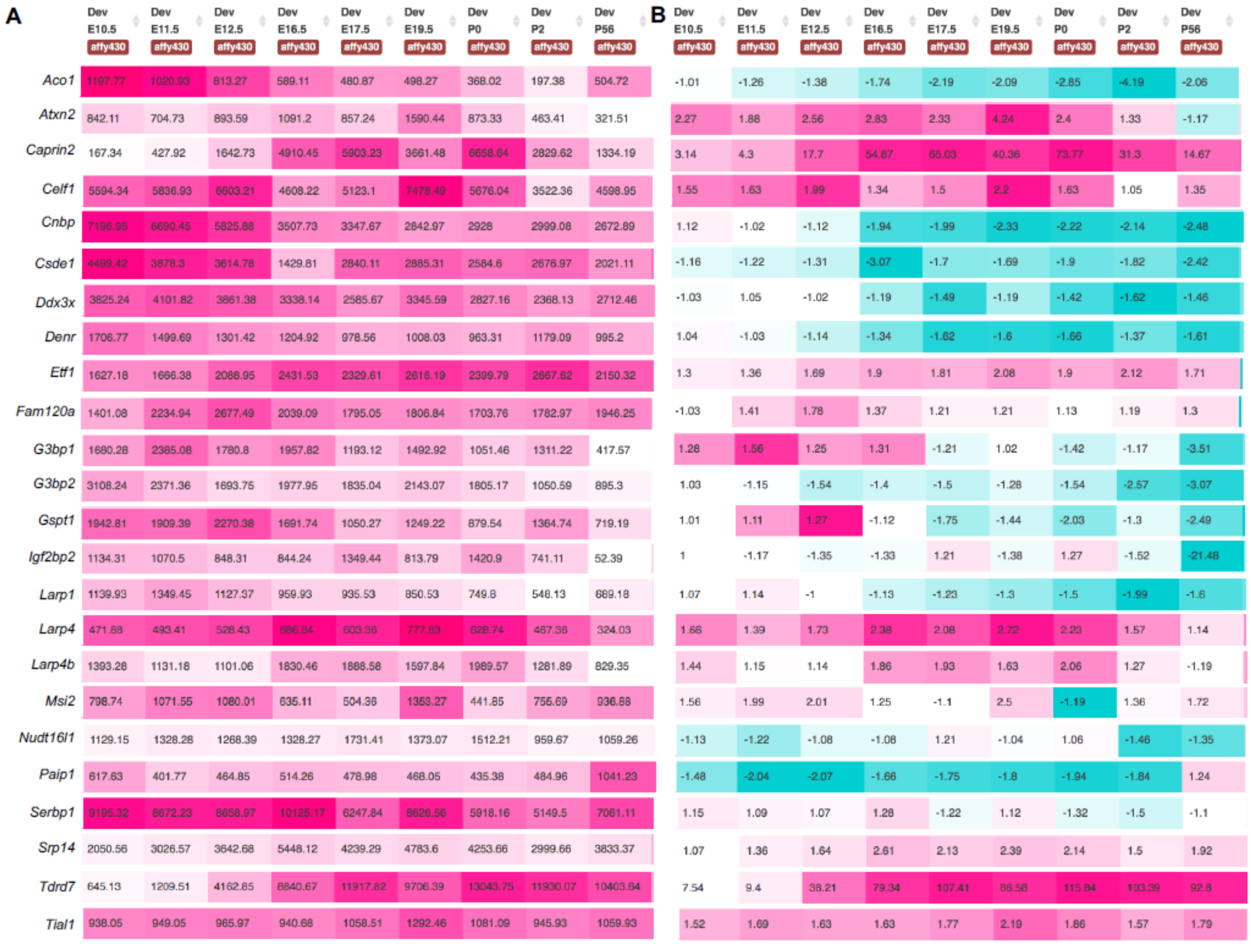

A

| | Dev E10.5 affy430 | Dev E11.5 affy430 | Dev E12.5 affy430 | Dev E16.5 affy430 | Dev E17.5 affy430 | Dev E19.5 affy430 | Dev P0 affy430 | Dev P2 affy430 | Dev P56 affy430 |
|---|---|---|---|---|---|---|---|---|---|
| *Aco1* | 1197.77 | 1020.93 | 813.27 | 589.11 | 480.87 | 498.27 | 368.02 | 197.38 | 504.72 |
| *Atxn2* | 842.11 | 704.73 | 893.59 | 1091.2 | 857.24 | 1590.44 | 873.33 | 463.41 | 321.51 |
| *Caprin2* | 167.34 | 427.92 | 1642.73 | 4910.45 | 5903.23 | 3661.48 | 6658.64 | 2829.62 | 1334.19 |
| *Celf1* | 5594.34 | 5836.93 | 6603.21 | 4608.22 | 5123.1 | 7478.49 | 5676.04 | 3522.36 | 4598.95 |
| *Cnbp* | 7196.95 | 6690.45 | 5825.88 | 3507.73 | 3347.67 | 2842.97 | 2928 | 2999.08 | 2672.89 |
| *Csde1* | 4499.42 | 3878.3 | 3614.78 | 1429.81 | 2840.11 | 2885.31 | 2584.6 | 2676.97 | 2021.11 |
| *Ddx3x* | 3825.24 | 4101.82 | 3861.38 | 3338.14 | 2585.67 | 3345.59 | 2827.16 | 2368.13 | 2712.46 |
| *Denr* | 1706.77 | 1499.69 | 1301.42 | 1204.92 | 978.56 | 1008.03 | 963.31 | 1179.09 | 995.2 |
| *Etf1* | 1627.18 | 1666.38 | 2088.95 | 2431.53 | 2329.61 | 2616.19 | 2399.79 | 2667.62 | 2150.32 |
| *Fam120a* | 1401.08 | 2234.94 | 2677.49 | 2039.09 | 1795.05 | 1806.84 | 1703.76 | 1782.97 | 1946.25 |
| *G3bp1* | 1680.28 | 2385.08 | 1780.8 | 1957.82 | 1193.12 | 1492.92 | 1051.46 | 1311.22 | 417.57 |
| *G3bp2* | 3108.24 | 2371.36 | 1693.75 | 1977.95 | 1835.04 | 2143.07 | 1805.17 | 1050.59 | 895.3 |
| *Gspt1* | 1942.81 | 1909.39 | 2270.38 | 1691.74 | 1050.27 | 1249.22 | 879.54 | 1364.74 | 719.19 |
| *Igf2bp2* | 1134.31 | 1070.5 | 848.31 | 844.24 | 1349.44 | 813.79 | 1420.9 | 741.11 | 52.39 |
| *Larp1* | 1139.93 | 1349.45 | 1127.37 | 959.93 | 935.53 | 850.53 | 749.8 | 548.13 | 689.18 |
| *Larp4* | 471.88 | 493.41 | 528.43 | 686.84 | 603.36 | 777.63 | 628.74 | 467.36 | 324.03 |
| *Larp4b* | 1393.28 | 1131.18 | 1101.06 | 1830.46 | 1888.58 | 1597.84 | 1989.57 | 1281.89 | 829.35 |
| *Msi2* | 798.74 | 1071.55 | 1080.01 | 635.11 | 504.36 | 1353.27 | 441.85 | 755.69 | 936.88 |
| *Nudt16l1* | 1129.15 | 1328.28 | 1268.39 | 1328.27 | 1731.41 | 1373.07 | 1512.21 | 959.67 | 1059.26 |
| *Paip1* | 617.63 | 401.77 | 464.85 | 514.26 | 478.98 | 468.05 | 435.38 | 484.96 | 1041.23 |
| *Serbp1* | 9195.32 | 8672.23 | 8658.97 | 10125.17 | 6247.84 | 8626.56 | 5918.16 | 5149.5 | 7061.11 |
| *Srp14* | 2050.56 | 3026.57 | 3642.68 | 5448.12 | 4239.29 | 4783.6 | 4253.66 | 2999.66 | 3833.37 |
| *Tdrd7* | 645.13 | 1209.51 | 4162.85 | 8840.67 | 11917.82 | 9706.39 | 13043.75 | 11930.07 | 10403.64 |
| *Tial1* | 938.05 | 949.05 | 965.97 | 940.68 | 1058.51 | 1292.46 | 1081.09 | 945.93 | 1059.93 |

B

| | Dev E10.5 affy430 | Dev E11.5 affy430 | Dev E12.5 affy430 | Dev E16.5 affy430 | Dev E17.5 affy430 | Dev E19.5 affy430 | Dev P0 affy430 | Dev P2 affy430 | Dev P56 affy430 |
|---|---|---|---|---|---|---|---|---|---|
| *Aco1* | -1.01 | -1.26 | -1.38 | -1.74 | -2.19 | -2.09 | -2.85 | -4.19 | -2.06 |
| *Atxn2* | 2.27 | 1.88 | 2.56 | 2.83 | 2.33 | 4.24 | 2.4 | 1.33 | -1.17 |
| *Caprin2* | 3.14 | 4.3 | 17.7 | 54.87 | 65.03 | 40.36 | 73.77 | 31.3 | 14.67 |
| *Celf1* | 1.55 | 1.63 | 1.99 | 1.34 | 1.5 | 2.2 | 1.63 | 1.05 | 1.35 |
| *Cnbp* | 1.12 | -1.02 | -1.12 | -1.94 | -1.99 | -2.33 | -2.22 | -2.14 | -2.48 |
| *Csde1* | -1.16 | -1.22 | -1.31 | -3.07 | -1.7 | -1.69 | -1.9 | -1.82 | -2.42 |
| *Ddx3x* | -1.03 | 1.05 | -1.02 | -1.19 | -1.49 | -1.19 | -1.42 | -1.62 | -1.46 |
| *Denr* | 1.04 | -1.03 | -1.14 | -1.34 | -1.62 | -1.6 | -1.66 | -1.37 | -1.61 |
| *Etf1* | 1.3 | 1.36 | 1.69 | 1.9 | 1.81 | 2.08 | 1.9 | 2.12 | 1.71 |
| *Fam120a* | -1.03 | 1.41 | 1.78 | 1.37 | 1.21 | 1.21 | 1.13 | 1.19 | 1.3 |
| *G3bp1* | 1.28 | 1.56 | 1.25 | 1.31 | -1.21 | 1.02 | -1.42 | -1.17 | -3.51 |
| *G3bp2* | 1.03 | -1.15 | -1.54 | -1.4 | -1.5 | -1.28 | -1.54 | -2.57 | -3.07 |
| *Gspt1* | 1.01 | 1.11 | 1.27 | -1.12 | -1.75 | -1.44 | -2.03 | -1.3 | -2.49 |
| *Igf2bp2* | 1 | -1.17 | -1.35 | -1.33 | 1.21 | -1.38 | 1.27 | -1.52 | -21.48 |
| *Larp1* | 1.07 | 1.14 | -1 | -1.13 | -1.23 | -1.3 | -1.5 | -1.99 | -1.6 |
| *Larp4* | 1.66 | 1.39 | 1.73 | 2.38 | 2.08 | 2.72 | 2.23 | 1.57 | 1.14 |
| *Larp4b* | 1.44 | 1.15 | 1.14 | 1.86 | 1.93 | 1.63 | 2.06 | 1.27 | -1.19 |
| *Msi2* | 1.56 | 1.99 | 2.01 | 1.25 | -1.1 | 2.5 | -1.19 | 1.36 | 1.72 |
| *Nudt16l1* | -1.13 | -1.22 | -1.08 | -1.08 | 1.21 | -1.04 | 1.06 | -1.46 | -1.35 |
| *Paip1* | -1.48 | -2.04 | -2.07 | -1.66 | -1.75 | -1.8 | -1.94 | -1.84 | 1.24 |
| *Serbp1* | 1.15 | 1.09 | 1.07 | 1.28 | -1.22 | 1.12 | -1.32 | -1.5 | -1.1 |
| *Srp14* | 1.07 | 1.36 | 1.64 | 2.61 | 2.13 | 2.39 | 2.14 | 1.5 | 1.92 |
| *Tdrd7* | 7.54 | 9.4 | 38.21 | 79.34 | 107.41 | 86.58 | 115.84 | 103.39 | 92.8 |
| *Tial1* | 1.52 | 1.69 | 1.63 | 1.63 | 1.77 | 2.19 | 1.86 | 1.57 | 1.79 |

#### Supplementary Table S1. smFISH hybridization probes.

| Target mRNA + Readout probes | Sequence |
|---|---|
| Cryaa_RO1_1 | ATACTGGAGCGACGCGTGATGTGGCCAGCATCCAAACCGGACTGGAATACTGGAGCGACGCGTGAT |
| Cryaa_RO1_2 | ATACTGGAGCGACGCGTGATTTGGGGCCAGAGAAGGTCAGCATGCCATACTGGAGCGACGCGTGAT |
| Cryaa_RO1_3 | ATACTGGAGCGACGCGTGATCCGCAGACAGGGAGCAGGAGAGGGCGATACTGGAGCGACGCGTGAT |
| Cryaa_RO1_4 | ATACTGGAGCGACGCGTGATTGGTCCACATTGGAAGGCAGACGGTAGCATACTGGAGCGACGCGTGAT |
| Cryaa_RO1_5 | ATACTGGAGCGACGCGTGATGCCCGAGTCCAGCACAGTGCGGAAGAATACTGGAGCGACGCGTGAT |

| | |
|---|---|
| Cryaa_RO1_6 | ATACTGGAGCGACGCGTGATTCAGGACGAGGGTGCAGAGCTGGGTTTCATACTGGAGCGACGCGTGAT |
| Cryaa_RO1_7 | ATACTGGAGCGACGCGTGATACCTTCGCCATGTAACTTGCCTTGACGATACTGGAGCGACGCGTGAT |
| Cryaa_RO1_8 | ATACTGGAGCGACGCGTGATGTTGTTCTTGGGGTTTCCAGCATGTGGTTGATACTGGAGCGACGCGTGAT |
| Cryaa_RO1_9 | ATACTGGAGCGACGCGTGATGACTGGCGGTAGTAGGGGCTGATGGTGATACTGGAGCGACGCGTGAT |
| Cryaa_RO1_10 | ATACTGGAGCGACGCGTGATAAGGCCCTCGCCGAAGAACTGGTCGAATACTGGAGCGACGCGTGAT |
| Cryaa_RO1_11 | ATACTGGAGCGACGCGTGATTAGAAGGGCCCCAGGGCACGCTTGAAATACTGGAGCGACGCGTGAT |
| Cryaa_RO1_12 | ATACTGGAGCGACGCGTGATCGGTGAAATTCACGGGAAATGTAGCCATGGTCATACTGGAGCGACGCGTGAT |
| Cryaa_RO1_13 | ATACTGGAGCGACGCGTGATCCTGCCTCTCGTTGTGTTTGCCGTGAATACTGGAGCGACGCGTGAT |
| Cryaa_RO1_14 | ATACTGGAGCGACGCGTGATCAAGAAGATGACAAACTTGTCCCGGTCAGATCATACTGGAGCGACGCGTGAT |
| Cryaa_RO1_15 | ATACTGGAGCGACGCGTGATGCATTACAAACCACATATGGGTCATGAGCTATACTGGAGCGACGCGTGAT |
| Cryaa_RO1_16 | ATACTGGAGCGACGCGTGATAGACAGGAAGGGCAGCAGGTCGTACTCAATACTGGAGCGACGCGTGAT |
| Cryaa_RO1_17 | ATACTGGAGCGACGCGTGATCTCCACAAAATCCTCCAGTACCTTCACATACTGGAGCGACGCGTGAT |
| Cryab_RO2_1 | GTTTGAAGATTCGACCTGGAAGAGAATCTGTCCTTCTCCAAACGCATCTGTTTGAAGATTCGACCTGGA |
| Cryab_RO2_2 | GTTTGAAGATTCGACCTGGACCGCAGGAAGGAGGGTGGCCGAAGGTAGGTTTGAAGATTCGACCTGGA |
| Cryab_RO2_3 | GTTTGAAGATTCGACCTGGAGGGGCTCAGGGAAGTGGCTGTTGAGAAGAGTTTGAAGATTCGACCTGGA |
| Cryab_RO2_4 | GTTTGAAGATTCGACCTGGATCAGACTCCAACAGGTGCTCTCCGAAGGTTTGAAGATTCGACCTGGA |
| Cryab_RO2_5 | GTTTGAAGATTCGACCTGGAAGGGGAAGAAGGGGCGCCGGATCCAGGTTTGAAGATTCGACCTGGA |
| Cryab_RO2_6 | GTTTGAAGATTCGACCTGGACTACTTCTTAGGGGCTGCGGCGACAGGTTTGAAGATTCGACCTGGA |
| Cryab_RO2_7 | GTTTGAAGATTCGACCTGGAGGCTTCTCTTCACGGGTGATGGGAATGGGTTTGAAGATTCGACCTGGA |
| Cryab_RO2_8 | GTTTGAAGATTCGACCTGGACTCAGGGCCAGACACCTGTTTCCTTGGTTTGAAGATTCGACCTGGA |
| Cryab_RO2_9 | GTTTGAAGATTCGACCTGGACTGGGATCCGGTACTTCCTGTGGAACTGTTTGAAGATTCGACCTGGA |
| Cryab_RO2_10 | GTTTGAAGATTCGACCTGGACTGGAGATGAAGCCATGTTCGTCCTGGCGTTTGAAGATTCGACCTGGA |
| Cryab_RO2_11 | GTTTGAAGATTCGACCTGGATCTTCGTGCTTGCCGTGGACCTCAATGTTTGAAGATTCGACCTGGA |

| | |
|---|---|
| Cryab_RO2_12 | GTTTGAAGATTCGACCTGGAGAGAGTCCGGTGTCAATCCAGCTGGGTGTTTGAAGATTCGACCTGGA |
| Cryab_RO2_13 | GTTTGAAGATTCGACCTGGACTGGTCGAAGAGGCGGCTTGGGGAGTGGTTTGAAGATTCGACCTGGA |
| Cryab_RO2_14 | GTTTGAAGATTCGACCTGGACCATTCACAGTGAGGACTCCATCAGATGAGTTTGAAGATTCGACCTGGA |
| Cryab_RO2_15 | GTTTGAAGATTCGACCTGGAGGGATGAAGTGATGGTGAGAGGATCCACATCGTTTGAAGATTCGACCTGGA |
| Cryab_RO2_16 | GTTTGAAGATTCGACCTGGACGTCCCCCAGAACCTTGACTTTGAGTTCGTTTGAAGATTCGACCTGGA |

#### Supplementary Table 2. RT-PCR primers

| Gene | Forward primer (5'-3') | Reverse primer (5'-3') |
|---|---|---|
| Cryaa | TCTGAGAGAGTGGCTTAGAG | GTGGCTGCAGTAAGTATGGG R |
| Cryab | CCGGAGGAACTCAAAGTCAA | AGCTTCAGCACTAGTCACAG |
| Crybb2 | TGAGAACCCCAACTTTACTG | AGCCACACTTTATTCTTCAC |
| Crygd | CGGCTCTCACAGGATCAGAC | CATGTCGTAGAGGACCCAGC |
| Cryga | TGGGTTTCAGCGACTCCATC | GTGGTAGCGCCTGTAGTCTC |
| β -actin | TGGCACCAGCACAATGAA | CTAAGTCATAGTCCGCCTAGAAGCA |

#### Supplementary Table 3. RBP motifs

| Protein | Motif |
|---|---|
| Pabpc1 | AGGACAAAACACT |
| Igf2bp3 | ACCAAATCATCAAG ;<br>ACGCATATCATCATCCAAC;<br>ACGCATATCATCATCCAAC |
| Ybx1 | CCATGTCATC;<br>GTACTGCAAGTATCCGATAGCGTAGCTG |
| Elavl1 | TGATATGAATGTGAGTAGATTGTGATAG;<br>TGATATGAATGTGAGTAGATTGTGATAG |
| Rbm24 | GTATAGCGTTGAGTATGAGTATAGGTTGAGTA |
| Ybx3 | TCGAAGTACCGATCAGTC |
| Carhsp1 | TACGTAGCGACTCTCTGTCCCTGAGTCTGAC;<br>TACGTAGCGACTCTCTGTCCCTGAGTCTGAC;<br>GCTCATGATCAGTGAGTTAGC |
| Pcbp1 | TGCACTCGATTACGTAGCTAGCGTACCTCTATCATAGC |
| Tardbp | GTAGGATGATGAGTTGTGTA; |

| | |
|---|---|
| | GTAGGATGATGAGTTGTGTA |
| Igf2bp1 (Zbp1) | |
